# Phylogenomics of *Brosimum* Sw. (Moraceae) and allied genera, including a revised subgeneric system

**DOI:** 10.1101/2020.05.15.098566

**Authors:** Elliot M. Gardner, Lauren Audi, Qian Zhang, Hervé Sauquet, Alexandre K. Monro, Nyree J.C. Zerega

## Abstract

We present a phylogenomic study of *Brosimum* and the allied genera *Trymatococcus* and *Helianthostylis*, with near-complete taxon sampling. Distributed from Mexico and the Greater Antilles to the Amazon, this clade contains the underutilized crop ramón (bread nut) (*Brosimum alicastrum*) as well as other species valued for timber or medicinal uses. Target enrichment for 333 genes produced a well-resolved phylogenetic tree and showed that *Trymatoccocus* and *Helianthostylis* are nested within *Brosimum*. We present a revised subgeneric classification of *Brosimum* based on phylogenetic and morphological considerations, including the reduction of *Trymatococcus* and *Helianthostylis* to subgenera. The monophyletic subgenera can be diagnosed based on stipule, pistillode, and cotyledon synapomorphies. Divergence date estimates suggest a Miocene origin for *Brosimum*, and ancestral area reconstruction indicated that all four subgenera originated and initially diversified in Amazonia before dispersing into other parts of South and Central America.

**Resumen:** Presentamos un estudio filogenómico del género *Brosimum* y sus aliados, *Trymatococcus* y *Helianthostylis*, y que incluye prácticamente todas las especies descritas. Su distribución va desde México y las Antillas Mayores hasta el Amazonas y comprende especies como el ramón (*B. alicastrum*), un cultivo infrautilizado, y otras especies empleadas como madera o en medicina. La secuenciación masiva dirigida de 333 marcadores nucleares de copia única permitió la reconstrucción de una filogenia bien resuelta, en la que se demuestra que *Trymatococcus* y *Helianthostylis* están anidados en *Brosimum*. Presentamos, por lo tanto, una clasificación revisada a nivel de especies, teniendo en cuenta los resultados moleculares y las características morfológicas, y donde *Trymatococcus* y *Helianthostylis* pasan a ser subgéneros de *Brosimum*. Estos subgéneros monofiléticos pueden ser identificados por caracteres de las estípulas y de los pistilodios.

## Introduction

The mulberry family (Moraceae) has approximately 1,100 species and 39 genera, with a worldwide distribution and a center of diversity in the tropics. It includes economically and ecologically important species such as breadfruit and jackfruit (*Artocarpus* J.R. Forst. & G. Forst.), mulberries (*Morus* L.) and figs (*Ficus* L.). Moraceae are characterized by latex in all parenchymatous tissue and tiny inconspicuous, unisexual flowers arranged in a wide variety of inflorescence forms, ranging from simple spikes to condensed heads, discs and the unique fig syconium.

While monophyly of Moraceae is not in doubt (Datwyler & Weiblen, 2004; Zerega & al., 2005; Zhang & al., 2011, 2019b), diverse morphology and widespread homoplasy within Moraceae has made the establishment of a robust and stable sub-familial taxonomy problematic (Corner, 1962; Berg, 1977, 2001; Rohwer, 1993; Clement & Weiblen, 2009; Gardner & al., 2020a). The most recent family-wide phylogenetic studies recognized seven tribes based on molecular and morphological evidence, with inflorescence morphology most closely reflecting the clades (Artocarpeae, Olmedieae, Dorstenieae, Ficeae, Maclureae, Moreae, and Parartocarpeae) (Clement & Weiblen, 2009; Zerega & Gardner, 2019; Gardner & al., 2020a). Of these, the Dorstenieae are particularly heterogenous, comprising species with both unisexual and bisexual inflorescences ranging from mulberry-like spikes (*Sloetia*) to condensed heads (*Brosimum*) or flattened discs (*Dorstenia*). This has led to conflicting views of phylogenetic relationships within the tribe (Berg, 2001; Clement & Weiblen, 2009; Zerega & al., 2010; Gardner & al., 2020a). A recent phylogenomic study of Dorstenieae, based on a target enrichment approach (Zhang & al., 2019a), confirmed the monophyly of Dorstenieae, which includes *Brosimum* together with fifteen other genera (Zerega & Gardner, 2019; Gardner & al., 2020a). These include members of the now-obsolete Brosimeae as well as others (*Bleekrodea, Broussonetia, Fatoua, Sloetia*, and *Sloetiopsis*) that were previously assigned to Moreae.

### Study system

*Brosimum* Sw. sensu Berg (Figure 1) comprises 15 neotropical species whose distribution extends from Mexico and the Greater Antilles to southern Brazil. The morphology and taxonomic history of the genus were most recently reviewed in detail by Berg (1972). Berg (1970) united the poorly differentiated genera *Brosimum, Galactodendrum, Ferolia* Aubl., and *Piratinera* Aubl. into *Brosimum* as currently circumscribed, based on inflorescence characters, maintaining *Ferolia* (Aubl.) C.C. Berg as a subgenus. The genera *Brosimum, Helianthostylis* (2 spp.) and *Trymatococcus* (2 spp.) comprised the tribe “Brosimeae” Tréc. (Berg, 1972), later included within Dorstenieae (Berg, 2001) but used here as an informal clade name. The species in these three genera are all latex-producing trees, native to habitats ranging from wet to seasonally dry forest. As is the case with most members of the Dorstenieae, inflorescences can be bisexual (but are not always so). “Brosimeae” inflorescence morphology is unique within Dorstenieae, typically consisting of a capitate inflorescence covered with many staminate (“male”) flowers and one (to several) central pistillate (“female”) flower immersed in the receptacle-like inflorescence axis, visible only by virtue of its exserted style (Figure 1C–D). The immature inflorescence is initially covered completely by peltate bracts, which may persist in fruit (Figure 1B,F). In fruit, the inflorescence axis becomes fleshy, surrounding the seed(s). Berg (1970, 1972) divided *Brosimum* into two subgenera: subgenus *Brosimum*, with non-amplexicaul stipules (Figure 1H) and more or less globose-capitate inflorescences, and subgenus *Ferolia*, with fully amplexicaul stipules (Figure 1J) and often with lobed inflorescences resembling small cauliflower heads. *Trymatococcus* and *Helianthostylis* closely resemble *Brosimum*, differing in the presence of pistillodes (always lacking in *Brosimum*), as well as inflorescence sexuality (always bisexual in *Trymatococcus* and always bisexual or pistillate in *Helianthostylis*), the presence of a well-developed staminate perianth (usually vestigial or lacking in *Brosimum*) and the number of stamens in the flower,. *Helianthostylis*, remarkable for its long pistillodes, protruding up to 2 cm from the staminate flowers, can also have unisexual staminate inflorescences.

**Figure 1.**
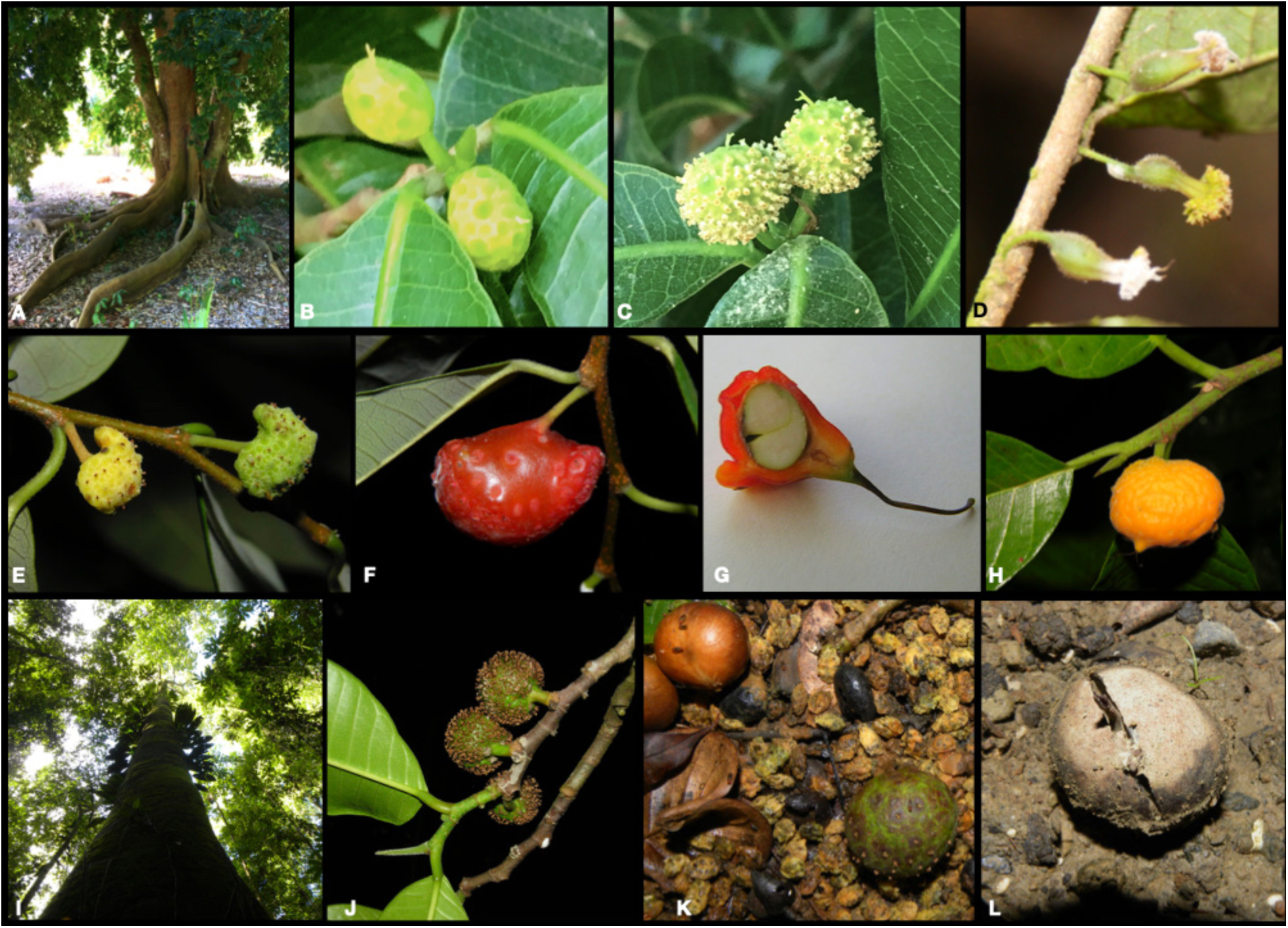
(A–C) *Brosimum alicastrum* Sw. buttresses, immature staminate inflorescences, and mature staminate inflorescences with central pistillate flower (the latter likely abortive); (D) *Trymatococcus amazonicus* inflorescences; (E–G) *Brosimum guianense* inflorescences, infructescene, and infructescence in section showing seed with unequal cotyledons typical of subg. *Brosimum*; (H) *Brosimum lactescens* shoot infructescence and lateral stipule typical of subg. *Brosimum*; (I–L) *Brosimum utile* bole, shoot with bisexual inflorescences showing amplexicaul stipule typical of subgenus *Ferolia*, seeds and infructescence showing persistent stamens, and germinating seed showing equal cotyledons typical of subg. *Ferolia*. Photos by E. Gardner (A–C), W. Milliken (D), Reinaldo Aguilar (E–F,H–L), and Alex Popovkin (G); photos E–L are reproduced under a CC BY-NC-SA 2.0 license, https://creativecommons.org/licenses/by-nc-sa/2.0/legalcode).

Several *Brosimum* species produce edible parts. The most well-known species, *B. alicastrum* (ramón, bread nut, Maya nut, Figure 1A–C) has a large nutritious seed; a traditional famine food, it has recently been promoted for its crop potential (Peters & Pardo-Tejeda, 1982; Gillespie & al., 2004; Lander & Monro, 2015). Additionally, several species have copious latex that is consumed as milk, including *B. utile* (palo de vaca, Figure 1I–L) (Berg, 1972). *Brosimum rubescens* (bloodwood) is a valuable timber tree, and *B. gaudichaudii* has been exploited by the pharmaceutical industry as a source of psoralens for the treatment of immunologic disorders (Palhares & al., 2007).

Previous phylogenetic work including *Brosimum* includes a two-locus family-level study (with five *Brosimum* species) based on nuclear ribosomal and chloroplast DNA (Zerega & al., 2005; Clement & Weiblen, 2009), a targeted study (12 *Brosimum* species) based on a single chloroplast locus (Silva, 2007), and a tribe-level study (10 *Brosimum* species) using a target enrichment approach (Zhang & al., 2019a). They all found that *Trymatoccocus* and *Helianthostylis* were nested within *Brosimum*. Some uncertainty remained, however, due to the small number of sequences in some cases and incomplete taxon sampling in others.

### Objectives

In order to resolve the relationships between these three genera, we employed target enrichment sequencing (HybSeq) to capture 333 genes previously developed for phylogenetic work in Moraceae (Gardner & al., 2016; Johnson & al., 2016), with nearly complete species-level sampling. The HybSeq method allows for efficient capture of hundreds of loci and is suitable for both fresh material and degraded DNA from herbarium material (Hart & al., 2016; Villaverde & al., 2018; Brewer & al., 2019), which comprises much of the material employed in this study. This work aims to test the monophyly of *Brosimum* and allied genera, determine synapomorphies for generic and subgeneric taxonomic levels, and revise *Brosimum* taxonomy accordingly. Additionally, we consider divergence date estimates and geographical distribution within the “Brosimeae,” testing whether lineages display a biogeographic pattern and reconstructing the ancestral range of the clade. While some species are widespread, others are restricted to the Amazon or the Guiana Shield. Recent studies in Moraceae have tended to reveal a strong correlation between biogeography and phylogeny.

## Materials and methods

### Sampling and DNA preparation

We sampled all 19 species of “Brosimeae” but were unable to obtain usable sequences from *B. glaziovii, B. melanopotamicum*, and *H. steyermarkii*, with final taxon sampling consisting of 16/19 species plus both subspecies of *B. alicastrum* (Table 1). Outgroups included at least one species from all other genera within *Dorstenieae* (12 additional genera), one sample per tribe for the remaining six Moraceae tribes, and *Trema orientale* (Cannabaceae). We prepared 22 new sequencing libraries for this study, combining those samples with reads from Zhang et al. (2019), Johnson et al. (2016), and GenBank to complete the outgroup sampling. All reads have been deposited in the National Center for Biotechnology Information (NCBI) Sequence Read Archive (BioProject PRJNA301299). Sample preparation followed the approach detailed in Hale et al. (2020). DNA was extracted from ca. 0.5 cm^2^ of leaf tissue (in almost all cases taken from a herbarium sheet) using a modified CTAB protocol (Doyle & Doyle, 1987), with overnight incubation for the initial lysis step as well as DNA precipitation. After assessing DNA fragment size on an agarose gel, samples with average fragment sizes more than 550 bp were sonicated to a mean insert size of 550 bp using a Covaris M220 (Covaris, Wobum, Massachussetts, USA). Libraries were prepared with the Illumina TruSeq Nano HT DNA Library Preparation Kit (Illumina, San Diego, California, USA) or the KAPA Hyper Prep (KAPA Biosystems, Cape Town, South Africa) following the manufacturer’s protocol, except that reactions were performed in one-third volumes to save reagent costs. Libraries were combined into pools of 6–24 and enriched for the 333 target genes (Gardner et al. 2016) with MYbaits custom probes (Arbor Biosciences, Ann Arbor, Michigan, USA) following the manufacturer’s protocol, with 14 PCR cycles. Products were sequenced on an Illumina MiSeq (2 x 300bp, version 3 chemistry) at the Field Museum of Natural History alongside samples for other studies in multiplexed runs of 30–70 samples each.

**Table 1.**
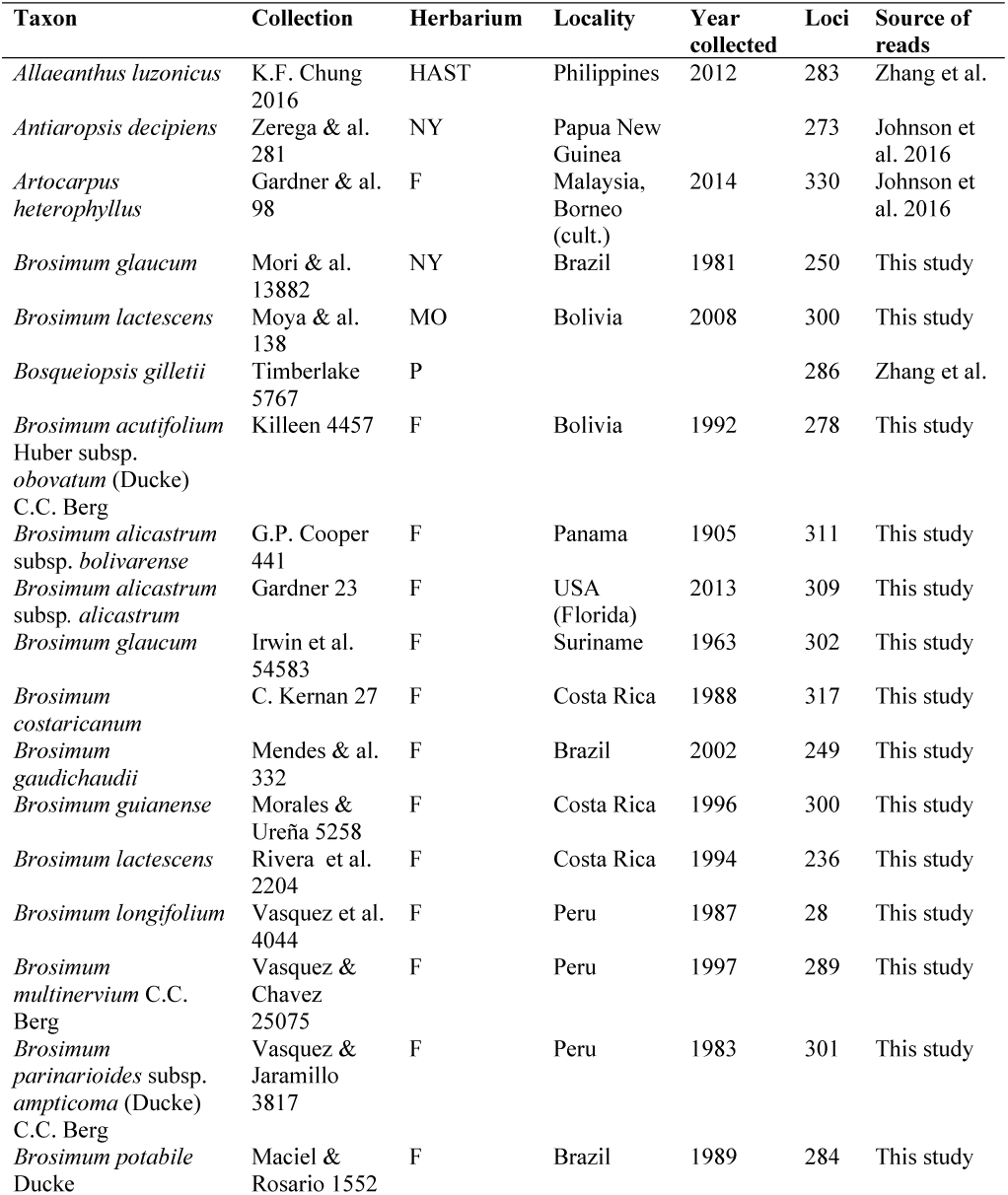
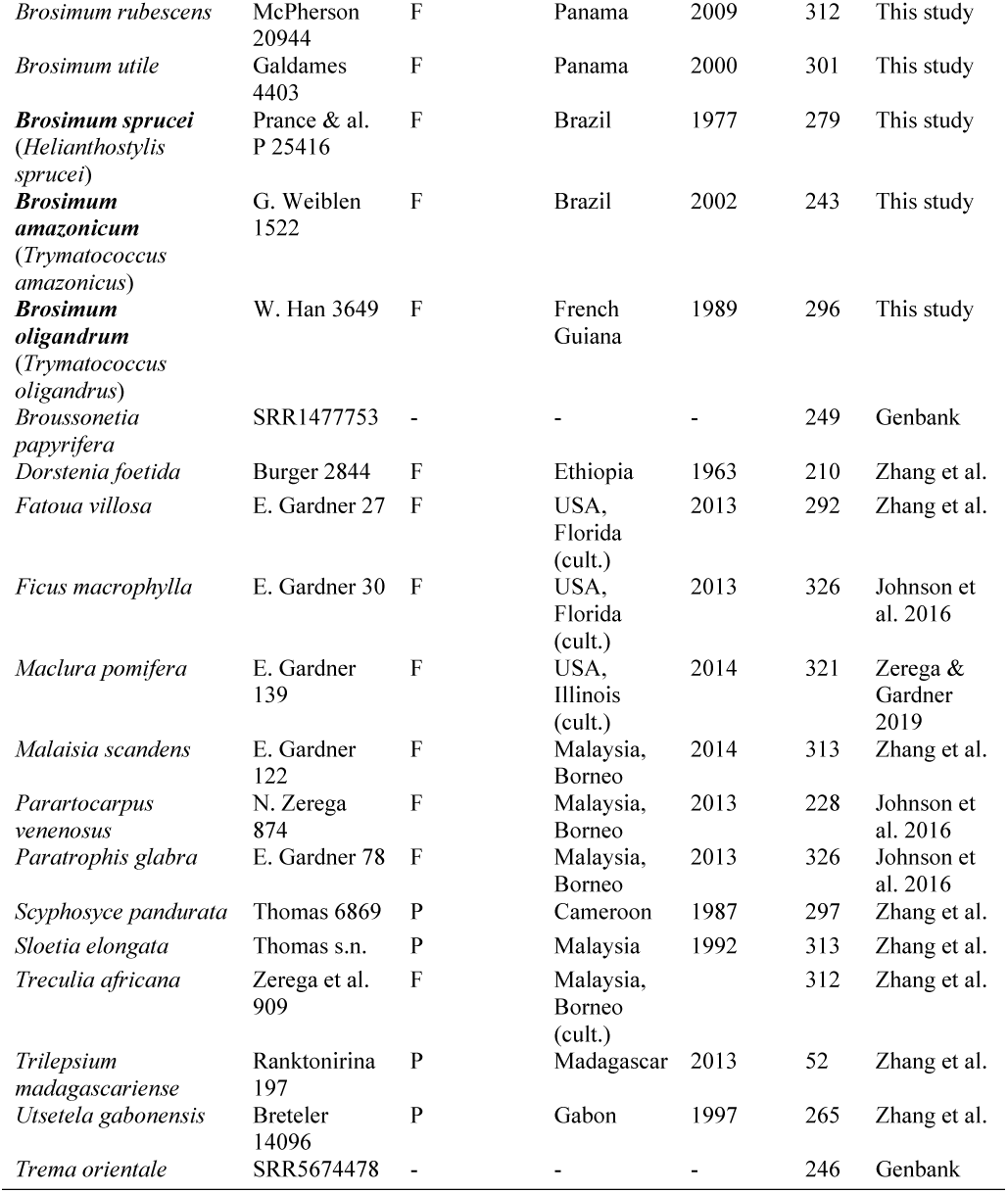
Accessions used in this study, including the number of loci recovered per sample, after filtering. Revised names appear in bold.

### Sequence assembly and phylogenetic analysis

We trimmed demultiplexed reads using Trimmomatic (with these parameters: “ILLUMINACLIP: TruSeq3-PE.fa:2:30:10 HEADCROP:3 LEADING:30 TRAILING:25 SLIDINGWINDOW:4:25 MINLEN:20”) (Bolger & al., 2014) and assembled them with HybPiper, which produces gene-by-gene, reference-guided, *de novo* assemblies. The default output of HybPiper is the predicted coding sequence for each gene (“exon” sequences). We used the HybPiper script “intronerate.py” to build “supercontig” sequences for each gene, consisting of exons as well as any assembled flanking non-coding sequences (intronic or intergenic).

For each target gene, “exon” sequences were filtered to remove sequences less than 25% of the average length for that gene. Filtered sequences were then aligned using MAFFT (Katoh & Standley 2013), and columns with more than 75% gaps were removed with Trimal (Capella-Gutiérrez et al. 2009). Single-gene phylogenies for each of 333 genes were estimated using RAxML under the GTRGAMMA model, with 500 rapid bootstrap replicates (Stamatakis 2006). After collapsing nodes with less than 30% bootstrap support (Zhang & al., 2017) using SumTrees (Sukumaran & Holder 2010), we used these gene trees to estimate a species tree with a coalescent-based approach implemented in ASTRAL-III (Mirarab & Warnow 2015, Zhang et al. 2018). Bootstrap (160 replicates) and local posterior probability (representing quartet support) were calculated for each node. A maximum likelihood tree was also calculated with RAxML based on a concatenated supermatrix of all 333 genes using a mixed-partition (one partition per gene) GTRGAMMA model, with 500 rapid bootstrap replicates. The analyses were then repeated with the “supercontig” sequences.

### Divergence time estimation

We time-calibrated the *Brosimum* phylogenetic tree with three fossils (for full details, see Zhang et al. 2019a) and two secondary calibrations. The fossil wood of *Artocarpoxylon deccanensis* Mehrotra, Prakash, and Bande (Mehrotra & al., 1984) was used as a minimum age constraint of 64.0 Ma for the stem node of *Artocarpus*, represented in our trees here by the split of *Artocarpus heterophyllus* and *Streblus glaber*. The fossil endocarps of *Broussonetia rugosa* Chandler (Chandler, 1961) were used to constrain to at least 33.9 Ma the stem node of *Broussonetia* s.s., represented in our trees by the most recent common ancestor of *Allaeanthus luzonicus, Malaisia scandens* and *Broussonetia papyrifera*. The fossil achenes of *Ficus* (*F. lucidus* Chandler) (Chandler, 1962) were used as a minimum age constraint of 56.0 Ma for the stem node of *Ficus*, represented in our trees by the split of *Ficus macrophylla* and *Antiaropsis decipiens*. Lastly, the estimated ages for the crown node of Moraceae (73.2-84.7 Ma) and the most recent common ancestor of Moraceae and Cannabaceae (81.7-93.3 Ma) obtained from a recent family-wide molecular dating (Zhang et al. 2019a) was used to secondarily calibrate the age of Moraceae and the root.

We estimated divergence times with both penalized likelihood (PL) and Bayesian relaxed clock approaches. PL was conducted in r8s v1.7 (Sanderson, 2003) using the tree from the RAxML analysis of the concatenated supermatrix with strict minimum (fossils) and maximum (root nodes) age constraints as described above. The best smoothing value was obtained using cross validation by testing 21 values of smoothing parameter scaling from 0.1 to 1000. The best value (i.e., with the lowest chi-square) of 2.6 was then applied as the smoothing parameter in divergence time estimation.

We used MCMCTree as implemented in the PAML v4.9 package (Yang, 2007) to estimate the divergence times with a Bayesian relaxed clock, using the topology from the best-known tree obtained from the RAxML analysis of the concatenated supermatrix. Two steps are needed for estimating divergence times by the approximate likelihood approach in MCMCTree. We first estimated the gradient and Hessian of branch length, and then used them to estimate the divergence times. A rough mean of several parameters were estimated by baseml in PAML v4.9 at first. Then we set the parameters in MCMCTree according to the results from baseml. We launched the Baysian analysis with a chain length of 22 million generations with the first 10% of the chain length discarded as burnin, sampling a total of 10,000 generations at a frequency of once every 2,000 generations. To confirm convergence, two independent runs with the same setting were conducted. To test the influence of the prior on the estimate, we used another prior setting (program defaults), followed the same process as described above, then ran the chain for 16.5 million generations, sampling a total of 10,000 generations at a frequency of once every 1,500 generations. An additional run was conducted for each setting but without data to confirm that the posterior was different from the prior. The convergence of the runs were checked using Tracer v1.7 (Rambaut & al., 2018). After confirming all the effective sample size (ESS) were over 100, we combined the results of the two independent runs for each prior setting. The entire dataset set was treated as a single partition to avoid extremely long running times. The substitution model used in MCMCTree was GTR with gamma.

### Ancestral state reconstructions

A morphological matrix of 16 categorical characters was constructed based on examination of herbarium specimens at the Royal Botanic Gardens Kew (K) and the published literature (Berg, 1972, 2001) (Appendix 1A). Characters included both vegetative and reproductive traits, including the stipule and pistillode characters used to difference the genera of the “Brosimeae” and the subgenera of *Brosimum* (Appendix 1A). We pruned the Bayesian time-calibrated tree to include only the Dorstenieae sensu stricto (*Brosimum, Helianthostylis, Trymatococcus, Bosqueiopsis, Dorstenia, Scyphosyce, Treculia, Trilepsium*, and *Utsetela*) and carried out ancestral state reconstruction under maximum-likelihood using the rayDISC function in corHMM 1.22, choosing the best fitting model between “ER,” “SYM,” and “ARD” based on AICc for each trait (Beaulieu & al., 2013). Manipulation and plotting of trees was carried out using ape and phytools in R (Revell, 2012; Paradis & Schliep, 2019; Team, 2019).

We assembled biogeographic matrices for the ingroup species based on the maps from Berg (1972) (Appendix 1B). For the first matrix, areas were based on the 10 Neotropical regions proposed by Antonelli et al. (2018), eight of which contained ingroup species: AMA (Amazonia), ATF (Atlantic Forests), AGL (Andean Grasslands), CAA (Caatinga), CEC (Cerrado and Chaco), DNO (Dry Northern South America), MES (Mesoamerica), and WIN (West Indies). Ranges were coded as present (1) or absent (0) and were compared to the “Brosimeae” records from that study, which originated from the Global Biodiversity Information Facility (GBIF). For the second matrix, we combined some areas that were adjacent and co-occurring and thus redundant and added the Guiana Shield as a separate area, as some ingroup species are concentrated there. The result contained four areas: Guiana Shield (including DNO), Amazonia (not including the Guiana Shield), Mesoamerica (including WIN), and Atlantic Forest (including CAA and CEC). For each matrix, we estimated ancestral ranges under the Dispersal-Extinction-Cladogenesis (DEC) model using BioGeoBEARS in R (Matzke, 2013), setting the maximum number of allowed areas to 4 (equal to the maximum number of areas currently occupied by any single species).

### Data availability

Raw reads were deposited in GenBank (BioProject PRJNA322184). Annotated workflow scripts detailing sequence assembly, alignment, and analysis parameters appear in Appendix 2.

## Results

### Phylogenetic analyses

Sequencing and assembly were successful for all taxa except *B. glaziovii, B. melanopotamicum*, and *Helianthostylis steyermarkii*; statistics appear in Table 1. The analyses were generally concordant. The inclusion of non-coding sequences did not affect the topology at all, and differences in the ingroup between the supermatrix and species-tree analyses were limited to a single rearrangement in the clade containing *B. paranarioides, B. longifolium, and B. potabile*; among outgroups, the positions of *Dorstenia, Scyphosyce*, and *Utsetela* also differed (Figure 2). In all analyses the “Brosimeae” as circumscribed by Berg (1972) were monophyletic, but, *Brosimum* was not, as *Helianthostylis* and *Trymatococcus* were nested within it. However, the subgenera were monophyletic. The sister group to the “Brosimeae” was *Treculia africana*.

**Figure 2.**
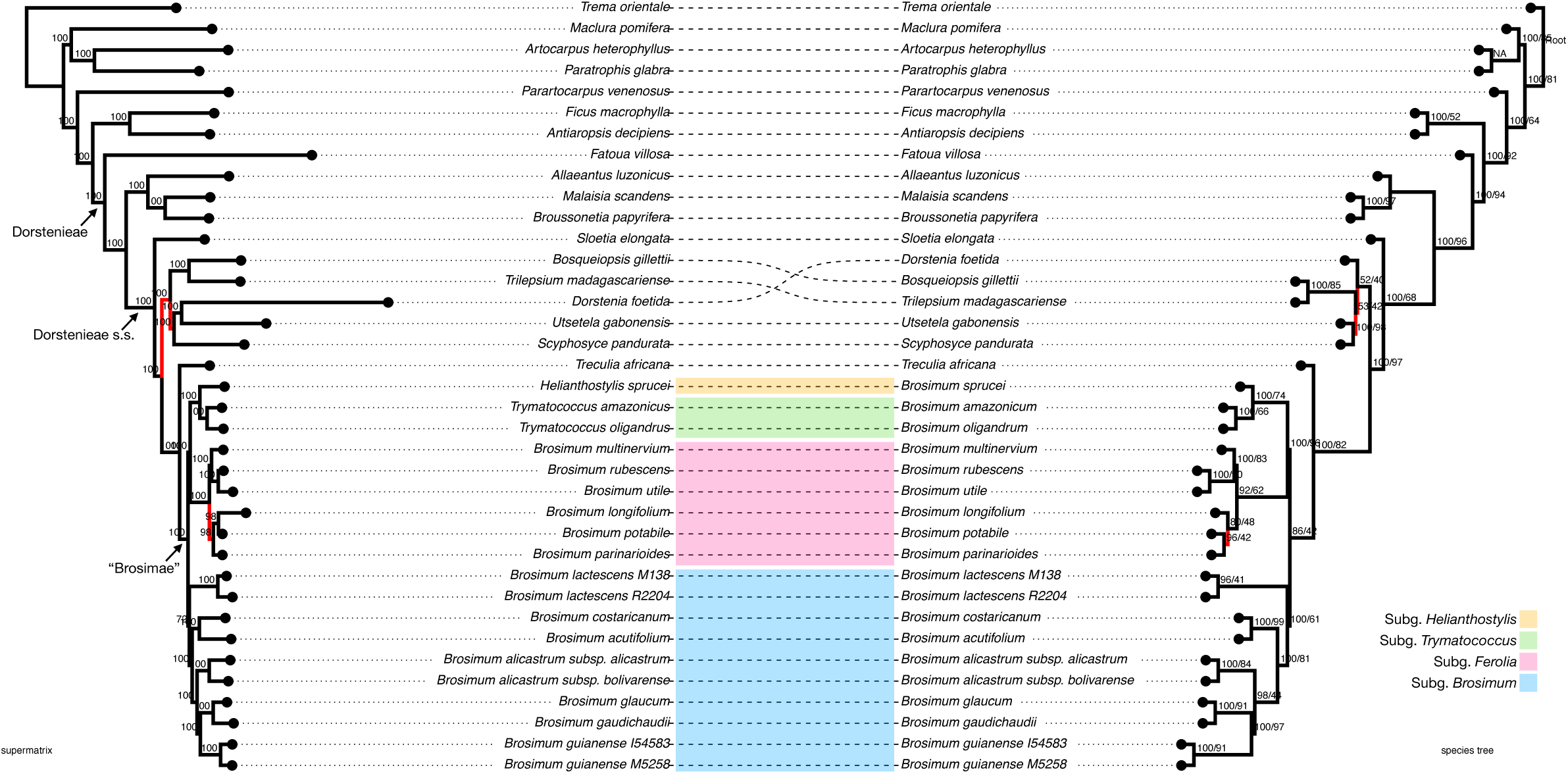
Left: maximum-likelihood tree based on a supermatrix of all sequences (“exon” dataset), showing bootstrap support, with previously accepted names. Right: ASTRAL species tree based on 333 gene trees (“exon” dataset), showing bootstrap/local posterior probability, with revised names. Disagreeing branches appear in red. The “supercontig” topologies were identical (and therefore not shown).

### Divergence time estimation

The stem- and crown-group ages of *Brosimum* s.l. were estimated to 22.09-35.35 Ma (early Miocene) and 18.49-29.62 Ma (Oligocene to early Miocene), respectively. Estimation from two different prior settings in MCMCTree showed similar results (Table 2, Figure S1). The ages estimated with a PL approach were consistent with the results from MCMCTree (Figure S2c) as well. Estimates from control runs without input data differed from results based on our dataset, suggesting the estimates were not determined by the priors.

**Table 2.**
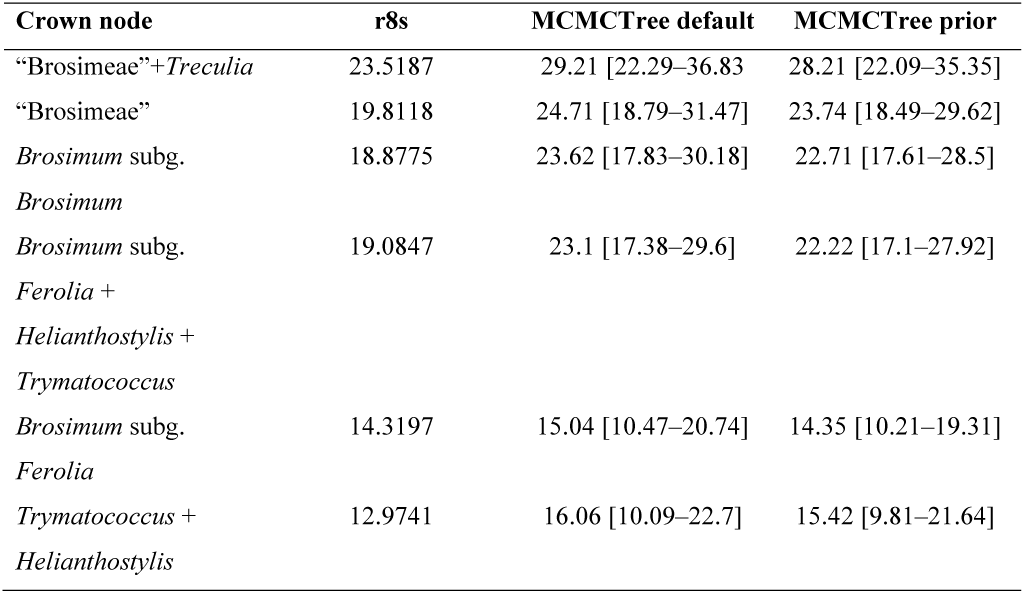
Estimated divergence times for select clades for all three analyses. 95% HPD values for the MCMCTree analyses appear in brackets.

### Ancestral state reconstructions

Trait mapping revealed several synapomorphic traits of taxonomic value in distinguishing the four sub-generic clades treated below (Figures 3, 4, S2). Amplexicaul stipules (always co-occurring with long stipules) are diagnostic of subgenus *Ferolia*. Pistillodes characterize only the *Trymatococcus*+*Helianthostylis* clade, but only *Trymatococcus* has protuberances on the pistillate inflorescences, and within that clade, only *Helianthostylis* has equal cotyledons (a character shared with *Ferolia*). Only subgenus *Brosimum* lack both amplexicaul stipules and pistillodes. Other traits appear to be homoplastic. For ancestral state reconstruction, model testing indicated that the “ER” model was preferred for all characters (Appendix 1C). revealed that the common ancestor of the ‘Brosimeae” most likely was a monoecious tree with non-amplexicaul stipules, bisexual, more or less globose inflorescences with peltate bracts and a single pistillate flower, staminate flowers lacking both well-developed perianth and a pistillode, and unequal cotyledons. The well-developed staminate perianth was likely regained three times, once each in *Trymatococcus+Helianthostylis, B. lactescens*, and *B. costaricanum*.

**Figure 3.**
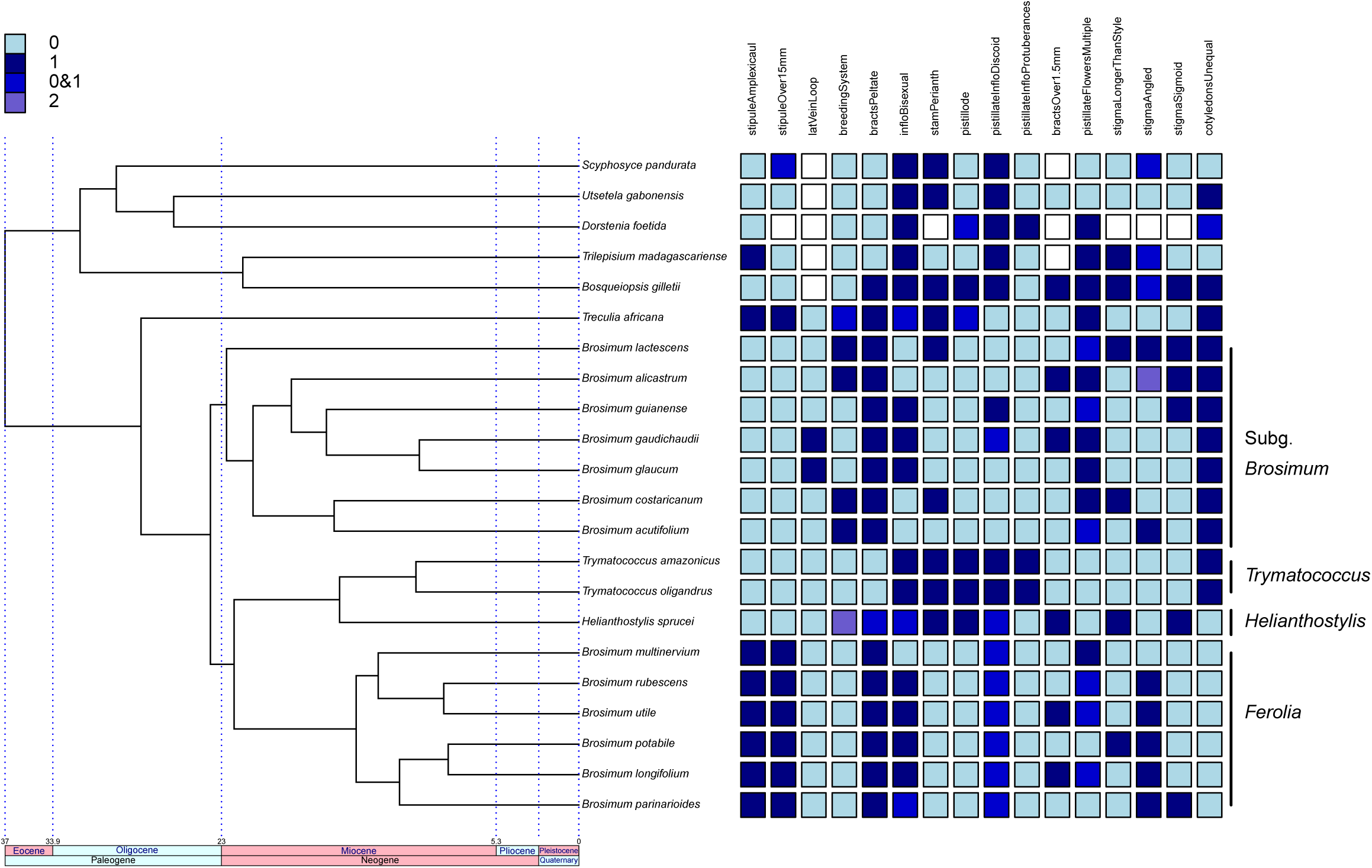
Sixteen characters mapped onto the pruned, time-calibrated phylogeny, with revised nomenclature, including subgenera. From left: (A) stipules fully amplexicaulous (0, 1); (B) longer stipules ≤ 15 mm (0), > 15 mm (1); (C) lateral veins loop-connected close (0) or far (1) from margin; (D) breeding system monoecious (0), dioecious (1), androdioecious (2); (E) interfloral bracts peltate (0/1); (F) inflorescences unisexual (0), bisexual (1); (G) Staminate perianth well developed (1) or vestigial / lacking (0); (H) pistillode absent (0), present (1); (I) pistillate inflorescence shape globose to ellipsoid (0) or turbinate, cylindrical, or hemispherical (1); (J) pistillate inflorescence surface with notable protuberences (1) or smooth (0); (K) Bracts greater than 1.5 mm (0, 1); (L) pistillate flowers solitary (0) or multiple (1); (M) stigma equal or shorter than style (0), longer than style (1); (N) stigma disposition angle from vertical: under 49 (0), 45-90 (1), over 90 (2); (O) stigma weakly curved (0), sigmoid; (P) cotyledons unequal (1) or not (0).

**Figure 4.**
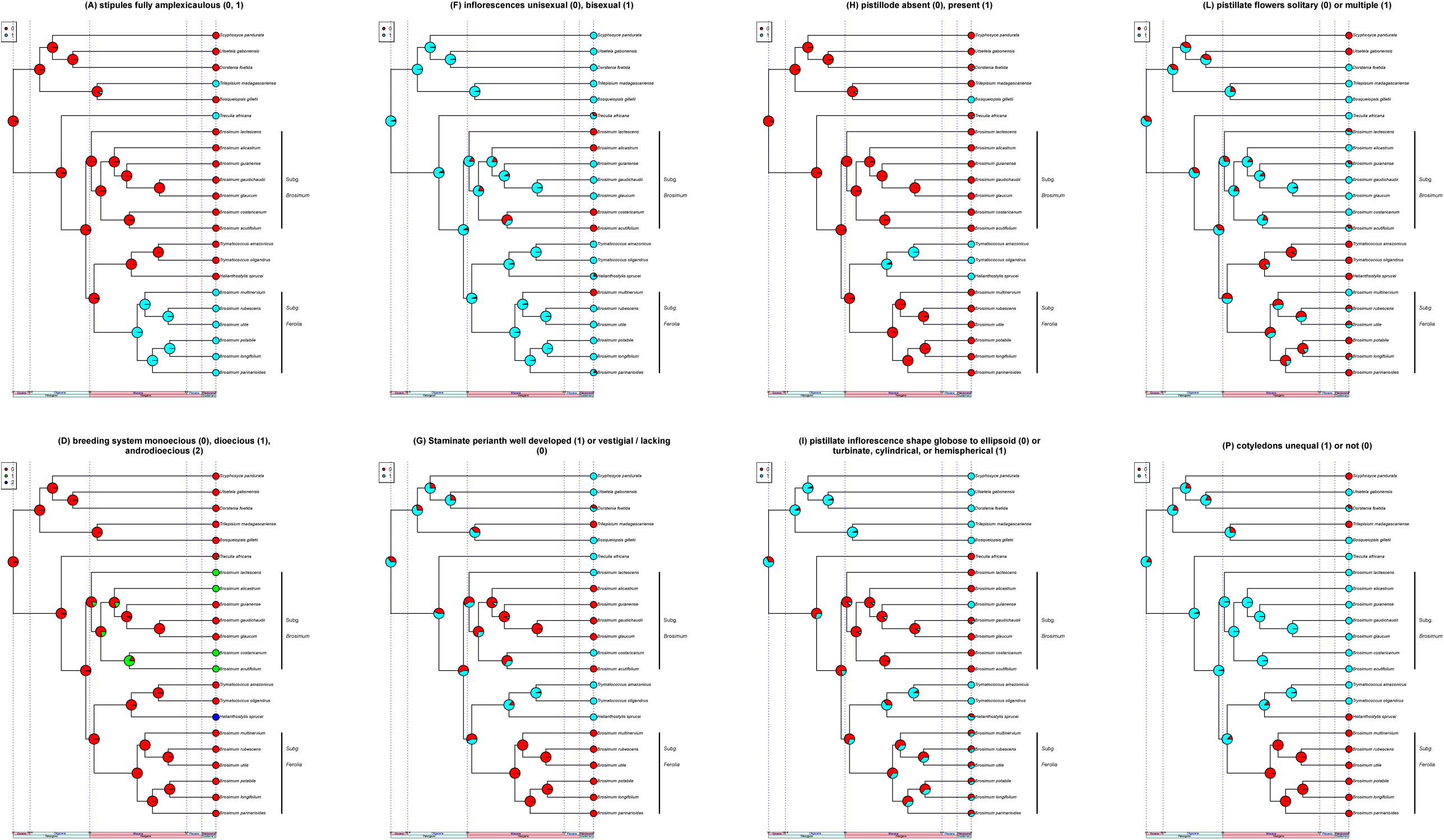
Maximum-likelihood ancestral reconstructions for eight select characters, labeled to align with the complete set of characters appearing in Figures 3 and S2. (A) stipules fully amplexicaulous (0, 1); (D) breeding system monoecious (0), dioecious (1), androdioecious (2); (F) inflorescences unisexual (0), bisexual (1); (G) Staminate perianth well developed (1) or vestigial / lacking (0); (H) pistillode absent (0), present (1); (I) pistillate inflorescence shape globose to ellipsoid (0) or turbinate, cylindrical, or hemispherical (1); (L) pistillate flowers solitary (0) or multiple (1); (P) cotyledons unequal (1) or not (0).

Both DEC analyses (Figures 5, S3) estimated Amazonia as the ancestral range of all four ingroup clades (subgenera *Brosimum* and *Ferolia, Trymatococcus*, and *Helianthostylis*), with dispersals into additional areas occurring beginning approximately 15 million years ago (8-area analysis: 1nL=-46.24084, nparams=2, d=0.007252254, e=1e-12, 4-area analysis: lnL=-40.27165, nparams=2, d=0.02100957, e=0.002998425). The 4-area analysis, which considered the Guiana Shield separately, estimated Amazonia+Guiana Shield as the ancestral range for *Trymatococcus* but estimated Amazonia not including the Guiana Shield as the ancestral range for the other three lineages, with subsequent dispersals to the Guiana Shield in the common ancestor of *B. potabile, B. longifolium*, and *B. parinaroides* (ca. 12 Ma) as well as even more recent dispersals in other individual species.

**Figure 5.**
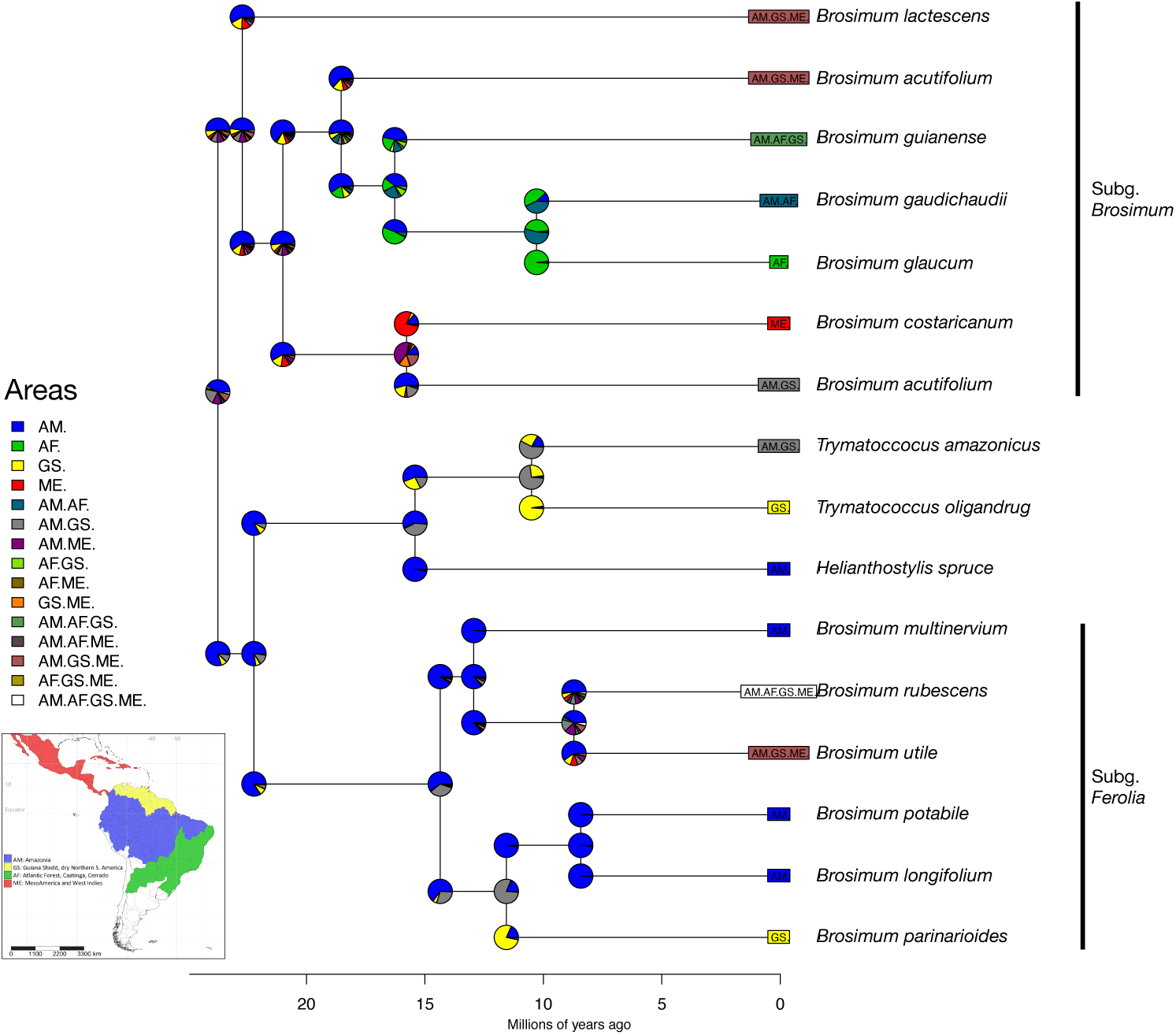
Results of DEC analysis using four simplified biogeographic areas. Pie charts indicate relative probabilities of different scenarios. Areas are AM (Amazonia, not including the Guiana Shield), GS (Guiana Shield and the adjacent Dry Northern South America), AF (Atlantic Forest and the adjacent Cerrado and Caatinga), and ME (Mesoamerica and West Indies).

## Discussion

### Phylogenetic relationships

The phylogenetic trees presented here, based on hundreds of genes with near-complete species sampling, were consistent with previous findings (Zerega et al. 2005, Clement and Weiblen 2009, Silva 2007, Zhang et al. 2019) that *Brosimum* is paraphyletic, encompassing *Trymatoccocus* and *Helianthostylis* (Figure 2). The morphological affinities between these genera, in particular the inflorescence architecture, led previous authors to treat them together as the “Brosimeae,” and notwithstanding the rank of *Trymatococcus* and *Helianthostylis*, our results match the higher divisions outlined by Berg in his morphology-based treatment of the “Brosimeae” (Berg 1972). Sister to the ingroup was the African genus *Treculia* (3 species native to Afrotropics), whose inflorescences, although larger, are also covered with peltate bracts that strongly resemble those of *Brosimum*. This sister placement of *Treculia* is consistent with previous studies (Zerega et al. 2005, Silva 2007, Clement and Weiblen 2009, Misiewicz and Zerega 2012, Zhang et al. 2019).

Our results highlight the value of the 333 loci employed here for resolving and clarifying the systematics of Moraceae. Phylogenomic studies based on these markers have so far produced robust phylogenetic hypotheses for Artocarpeae (Gardner & al., 2020b), Dorstenieae (Zhang & al., 2019a), Moreae (Gardner & al., 2020a), and Parartocarpeae (Zerega & Gardner, 2019) and therefore show promise in resolving longstanding taxonomic problems in the family.

### Divergence times and historical biogeography

Divergence time estimates (Table 2, Figure S1) were consistent, but slightly younger than the results obtained in a Dorstenieae-wide analysis in which stem- and crown-group ages for *Brosimum* s.l. were estimated between 19.4–42.9 Ma and 14.7-31.7 Ma, respectively (Zhang et al. 2019b). These differences may have resulted from sparser sampling outside the “Brosimeae” in the present study. Africa and South America were last connected approximately 105 million years ago (McLoughlin 2001), but *Brosimum* seed dispersers include birds and bats (Monterrubio-Rico & al., 2009; Poelchau & Hamrick, 2012). The origin of *Brosimum* in the Neotropics might therefore be explained by long-distance dispersal from Africa (Zhang et al. 2019b). In assembling the area matrix, the distributions we coded based on the *Flora Neotropica* (Berg 1972) maps did not materially differ from the distributions coded from the cleaned GBIF data from Antonelli et al. (2018) (Appendix 1B). Estimates of ancestral ranges indicate that all four major lineages of “Brosimeae” originated in Amazonia and initially diversified there, first dispersing to other areas, such as Northern South America, and Central America in the middle Miocene (subgenus *Brosimum*) and to the Guiana Shield in the late Miocene (subgenus *Ferolia, Trymatococcus*, and *Helianthostylis*) (Figures 5, S3). These findings are consistent with overall patterns of Neotropical diversification, which are overwhelmingly centered in Amazonia (Antonelli & al., 2018).

### Morphological evolution

Stipule amplexicauly and equal cotelydons are derived states within the “Brosimeae” and comprise a synapomorphy for the subgenus *Ferolia* (Figure 4). In “Brosimeae” stipule amplexicaulity always co-occurred with longer stipules, perhaps due to geometric constraints because an amplexicaul stipule clasps the full circumference of the stem as well as the developing leaf (Figure 1J). In the broader Dorstenieae s.s., the trait is homoplastic and likely evolved independently twice outside our ingroup, in *Treculia*—sister to the “Brosimeae”— and in *Trilepsium*. The role of amplexicauly relative to lateral stipules is unclear. They may facilitate the production of large leaves through a protective and or, photosynthetic role and the clade comprising species with amplexicaul stipules does comprise the larger leaved ingroup taxa. The role of unequal cotyledons (Figure 1G) is unclear. They do not correlate with seed size so may instead relate to seedling establishment. As with amplexicaul stipules, equal cotyledons (are homoplastic within the Dorstenieae s.s., present also in *Trilepsium, Scyphosyce*, and *Dorstenia* (p.p.).

The presence of a pistillode is a synapomorphy for the *Trymatococcus+Helianthostylis* clade and is apparently derived within the Dorstenieae s.s., otherwise appearing only in *Bosqueiopsis* (Figure 4). The presence of a well-developed pistillode in Moraceae may facilitate the positioning of inflexed stamens that ballistically release pollen at anthesis (Corner, 1962), and it is therefore not surprising that most members of the Dorstenieae s.s.—which never have ballistic pollen release—would lack a pistillode. In that case, there is likely another function for the pistillode in *Trymatococus* and *Helianthostylis*, and it is possible that at least in the latter, the long showy pistillodes may relate to pollinator attraction. However, little is known about pollination in the Dorstenieae (Berg, 2001).

Berg (1972) separated the three genera of his “Brosimeae” based on a suite of characters (stipule amplexicaulity, pistillode, inflorescence sexuality, staminate perianth, and stamen number) that align with our results, requiring only a change of rank for *Trymatococcus* and *Helianthostylis*, highlighting the durability of his careful morphological studies. The stipule and pistillode characters may be readily observed in fertile specimens. However, the other characters, while informative in studying broader patters of evolution, are less reliable for diagnosing individual specimens. Inflorescence sexuality can vary within a species, sometimes within the same individual (Peters, 1991), leading Berg to cast doubt on its reliability as a diagnostic trait (Berg 1972). The presence of a well-developed staminate perianth, while readily observable, was recovered as homoplastic and is thus diagnostic only in combination with other characters. In contrast to other tribes in Moraceae, stamen number is remarkably variable within the genera of the Dorstenieae. Although *Trymatococcus* and *Helianthostylis* never have fewer than three stamens, there otherwise exists some overlap in stamen number between the subgenera treated here. We therefore find that the most reliable and phylogenetically-consistent diagnostic characters for distinguishing these clades are stipule amplexicaulity, the presence of a pistillode, and equal/unequal cotyledons (although the latter can be difficult to determine from herbarium specimens) (Figure 4).

### Taxonomic revisions

Based on these analyses, we reduce *Trymatoccocus* and *Helianthostylis* to subgenera of *Brosimum.* This expanded genus unites all the woody members of Dorstenieae species in the New World, all based on a shared inflorescence plan. We include *B. multinervium* in subgenus *Ferolia*; although it was described after Berg’s treatment of that subgenus in *Flora Neotropica Monograph 7*, the fully amplexicaul stipules and its phylogenetic position leave no doubt as to its correct placement. Likewise, we maintain Berg’s (1972) placement of the taxa we were unable to include in our phylogenetic analyses based on unambiguous morphological synapomorphies. The fully-amplexicaul stipules of *B. melanopotamicum* leave no doubt about its proper placement in subgenus *Ferolia*, where it also appeared, with strong support, in a previous single-locus phylogenetic study (Silva 2007). The lateral stipules and unequal cotyledons of *B. glaziovii* place it within subgenus *Brosimum*, even in the absence of sequence data. Finally, the non-amplexicaul stipules, equal cotyledons, and well-developed pistillodes of *H. steyermarkii* confirm its proper placement in subgenus *Helianthostylis*. Below follows an updated phylogenetic classification and taxonomic summary for the expanded genus *Brosimum*. A short-form description and summary of the key characters defining each group is provided. For complete taxonomic histories, synonymies, and descriptions, the reader should refer to C.C. Berg’s comprehensive treatments of *Brosimum, Trymatoccocus*, and *Helianthostylis* in *Flora Neotropica* monographs 7 and 83 (Berg, 1972, 2001).

**B****rosimum** Swartz, Prod. Veg. Ind. Occ. 12 (1788), nom. cons. Type: *B. alicastrum* O. Swartz (typ. cons.)

*Alicastrum* P. Browne, Civil and Natural History of Jamaica 372 (1765), nom. rejic.

*Piratinera* Aublet, Pl. Gui. 2: 888 (1775), nom. rejic.

*Ferolia* Aublet, Pl. Gui. Suppl. 7 (1775), nom. rejic.

*Galactodendrum* Kunth in Humboldt & Bonplant, Voyage, Rel. Hist. 2: 108 (1819).

*Trymatoccocus* Poepp. & Endl., Nov. Gen. 2: 30 (1838).

*Helianthostylis* Baillon, Adansonia 11: 299 (1875).

*Brosimopsis* S. Moore, Trans. Linn. Soc. II. 4: 473 (1895)

Monoecious or dioecious trees. ***Leaves*** distichous, pinnately veined. Stipules free or connate, lateral or fully amplexicaul. ***Inflorescences*** unisexual or bisexual, axillary, solitary or paired, globose, hemispherical, cylindrical, or turbinate, interfloral bracts usually peltate. Staminate flowers few to many, perianth well developed or lacking/vestigial, stamens 1–4, straight in bud, pistillode present or absent. Pistillate flowers one to several, immersed in the center of the inflorescence axis with the style exserted. ***Fruits*** adnate to the enlarged and fleshy inflorescence axis, to at least 2 cm in diameter; cotyledons equal or unequal.

19 species, restricted to the Neotropics; the only woody Dorstenieae in that region (Figure 6).

**Figure 6.**
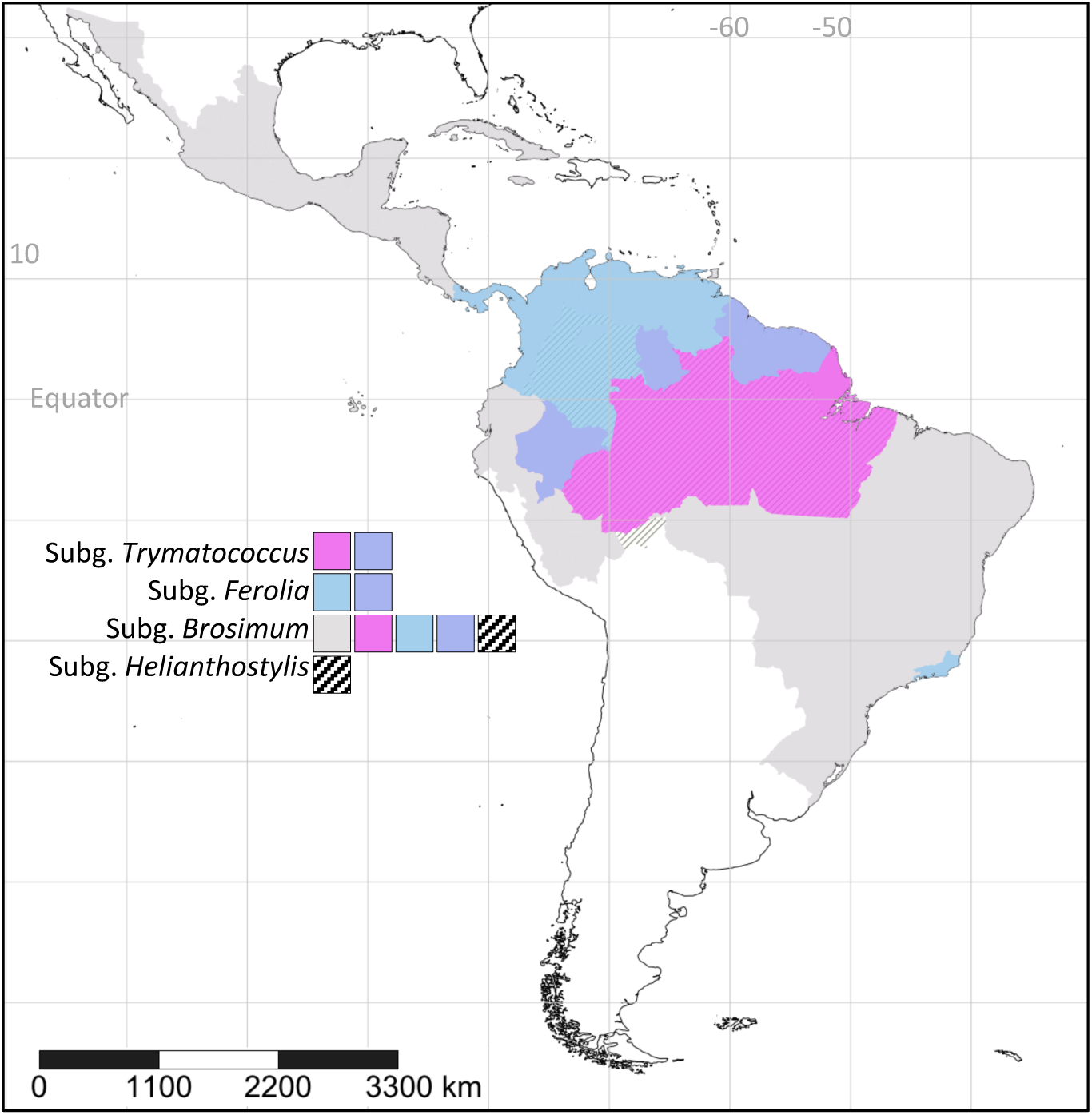
Distributions of the four subgenera of *Brosimum*, following the revisions outlined here.

#### Key to the subgenera

1. Pistillode absent:
  2. Stipules non-amplexicaul, cotyledons unequal – subgenus *Brosimum*
  2. Stipules fully amplexicaul, cotyledons equal – subgenus *Ferolia*

1. Pistillode present:
  3. Pistillode minute; cotyledons unequal, inflorescences with protuberances – subgenus *Trymatoccocus*
  3. Pistillode usually well-developed, cotyledons equal, inflorescences lacking protuberances – subgenus *Helianthostylis*

### Subgenus *Brosimum*

Monoecious or dioecious trees, stipules lateral, inflorescences unisexual or bisexual, usually ±globose, staminate perianth usually not well developed, pistillode absent, cotyledons unequal.

Distribution: Mexico and the Greater Antilles to northern South America and Amazon basin. (Colombia, Venezuela, Ecuador, Peru, Brazil, Bolivia, French Guiana, Costa Rica, Panama, Mexico, Guatemala, Nicaragua, Paraguay) (Figure 6).

Species (eight): *B. alicastrum* Sw., *B. acutifolium* Huber, *B. costaricanum* Liebm., *B. gaudichaudii* Trécul, *B. glaucum* Taub., *B. glaziovii* Taub., B. *guianense* (Aubl.) Huber ex Ducke, *B. lactescens* (S.Moore) C.C.Berg

### Subgenus *Ferolia* (Aublet) C.C. Berg

Monoecious trees, stipules amplexicaul, inflorescences usually bisexual, globose to, hemispherical, cylindrical, or turbinate, sometimes irregularly lobed, staminate perianth not well developed, pistillode absent, cotyledons equal.

Distribution: Members of this subgenus are largely concentrated in the northern part of the South American continent distinctly associated with the Guiana Shield (Panama, Colombia, Venezuela, Ecuador, Peru, Brazil, Bolivia, Guyana, French Guiana) (Figure 6).

Species (seven): *B. rubescens* Taub., *B. melanopotamicum* C.C. Berg, *B. utile* (Kunth) Oken, *B. longifolium* Ducke, *B. multinervium* C.C. Berg, *B. potabile* Ducke, *B. parinarioides* Ducke

**Subgenus *Trymatoccocus* (Poepp. & Endl.) E.M. Gardner & N.J.C. Zerega, stat. nov.** — based on *Trymatoccocus* Poepp. & Endl., Nov. Gen. 2: 30 (1838).

Monoecious trees, stipules lateral, inflorescences bisexual, turbinate, with protuberances, staminate perianth well developed, pistillode present but minute, cotyledons unequal.

Distribution: Upper Amazon basin to the Guianas (Colombia, Venezuela, Guyana, French Guiana, Ecuador, Peru, Brazil) (Figure 6).

> Species (two):
>
> ***Brosimum amazonicum* (Poepp. & Endl.) E.M. Gardner & N.J.C. Zerega, comb. nov** — based on *Trymatoccocus oligandrus* Peopp. & Endl., Nov. Gen. Sp. Pl. 2: 30 (1838).
>
> ***Brosimum oligandrum* (Benoist) E.M. Gardner & N.J.C. Zerega, comb. nov.** — based on *Lanessania oligandra* Benoist, Bull. Mus. Natl. Hist. Nat. 27: 199 (1921).

**Subgenus *Helianthostylis* (Baillon) E.M. Gardner & N.J.C. Zerega, stat. nov.** — based on *Helianthostylis* Baillon, Adansonia 11: 299 (1875).

Monoecious or androdioecious trees, stipules lateral, inflorescences bisexual or staminate, globose to turbinate, staminate perianth well developed, pistillode well developed and often showy, cotyledons equal.

Distribution: Amazon Basin (Brazil, Colombia, Ecuador, French Guiana, Guyana, Peru, Venezuela) (Figure 6).

> Species (two):
>
> ***Brosimum sprucei* (Baillon) E.M. Gardner & N.J.C. Zerega, comb. nov** — based on *Helianthostylis sprucei* Baill., Adansonia 11: 299 (1875).
>
> ***Brosimum steyermarkii* (C.C. Berg) E.M. Gardner & N.J.C. Zerega, comb. nov** — based on *Helianthostylis steyermarkii* C.C.Berg, Acta Bot. Neerl. 21: 99, fig. 1 (1972).

## Conclusion

The analyses and revisions presented above result in a well-defined and monophyletic *Brosimum* with four diagnosable subgenera, providing a framework for further work on this understudied genus, including phylogenetic and biogeographic investigations at the subspecific level.

## Supporting information

Appendices: 1A. Character matrix; 1B Area matrix; 1C Anc. recons. model testing. Appendix 2: Analysis workflow scripts

## Author contributions

EMG led the writing of the manuscript, did field, lab, and herbarium work, and conducted all analyses except the divergence time estimation. LA did herbarium sampling and lab work for the ingroup. QZ did herbarium sampling and lab work for the outgroups and conducted the divergence time analyses. HS acquired funding for outgroup sequencing and supervised overall work on Dorstenieae. AM provided overall guidance on *Brosimum* was responsible for collecting the morphological data including reviewing herbarium specimens and assembling the character matrix. NJCZ initiated the project, provided overall supervision, and acquired funding for ingroup sequencing. All authors commented on and contributed to the manuscript.

## Acknowledgements

This work was supported by the United States National Science Foundation (DEB awards 0919119 and 1501373 and DBI award 1711391), a China Scholarship Council (CSC) PhD grant (grant No. 201506140077) to Q.Z., and a research grant from the International Association for Plant Taxonomy (IAPT) to Q.Z. We thank the Pritzker Laboratory for Molecular Systematics at the Field Museum of Natural History (K. Feldheim) for the use of sequencing facilities, the Kampong (National Tropical Botanical Garden) for access to living collections, Louis Ronse De Craene (E) for advice on the use of anatomical terms, L. Pokorny for proofreading the Spanish abstract, and the following herbaria for access to specimens for examination and DNA extraction: F (C. Niezgoda), IBSC (Chung K.F.), MO (M. Merello), NY (M. Pace), and P (C. Sarthou).

