## Supplemental figures 1-3 for "Phylogenomics of *Brosimum* Sw. (Moraceae) and allied genera, including a revised subgeneric system"

Figure S1. Results of divergence time estimation by a) MCMCTree prior setting; b) MCMCTree default setting; c) r8s.

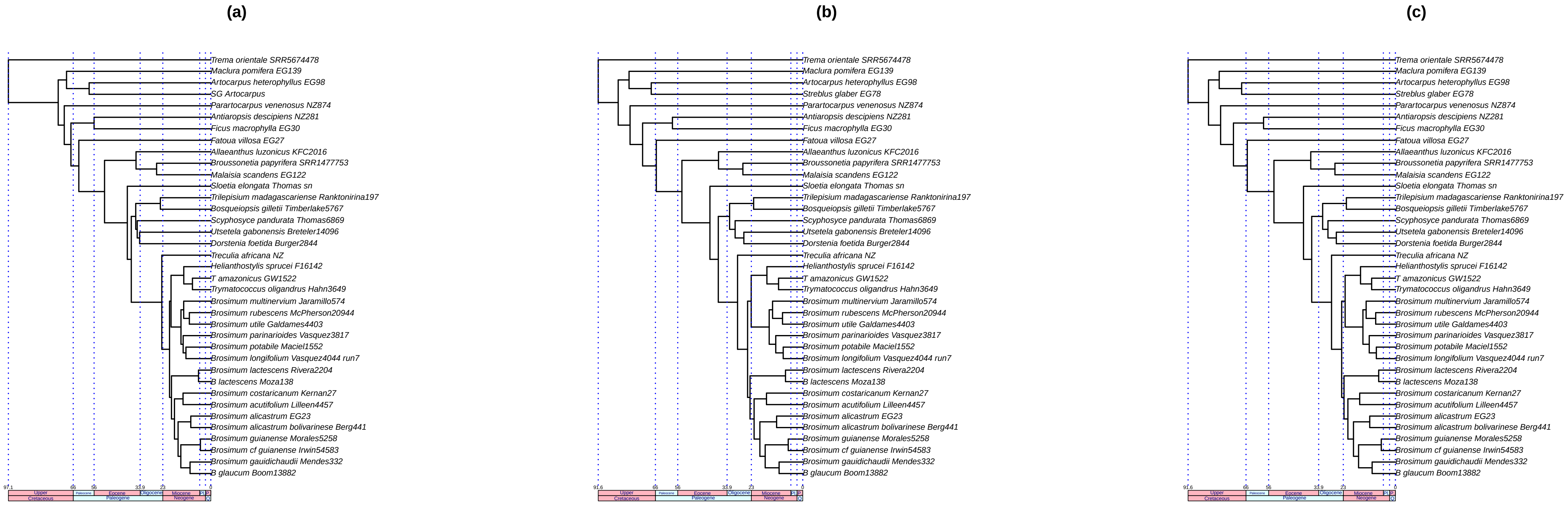
(a)
Trema orientale SRR5674478
Maclura pomifera EG139
Artocarpus heterophyllus EG98
SG Artocarpus
Parartocarpus venenosus NZ874
Antiaropsis descipiens NZ281
Ficus macrophylla EG30
Fatoua villosa EG27
Allaeanthus luzonicus KFC2016
Broussonetia papyrifera SRR1477753
Malaisia scandens EG122
Sloetia elongata Thomas sn
Trilepisium madagascariense Ranktonirina197
Bosqueiopsis gilletii Timberlake5767
Scyphosyce pandurata Thomas6869
Utsetela gabonensis Breteler14096
Dorstenia foetida Burger2844
Treculia africana NZ
Helianthostylis sprucei F16142
T amazonicus GW1522
Trymatococcus oligandrus Hahn3649
Brosimum multinervium Jaramillo574
Brosimum rubescens McPherson20944
Brosimum utile Galdames4403
Brosimum parinarioides Vasquez3817
Brosimum potabile Maciel1552
Brosimum longifolium Vasquez4044 run7
Brosimum lactescens Rivera2204
B lactescens Moza138
Brosimum costaricanum Kernan27
Brosimum acutifolium Lilleen4457
Brosimum alicastrum EG23
Brosimum alicastrum bolivarinese Berg441
Brosimum guianense Morales5258
Brosimum cf guianense Irwin54583
Brosimum gauidichaudii Mendes332
B glaucum Boom13882
97.1
66
56
33.9
23
0
Upper
Cretaceous
Paleocene
Eocene
Oligocene
Paleogene
Miocene
Neogene
Pl
P
Q
(b)
Trema orientale SRR5674478
Maclura pomifera EG139
Artocarpus heterophyllus EG98
Streblus glaber EG78
Parartocarpus venenosus NZ874
Antiaropsis descipiens NZ281
Ficus macrophylla EG30
Fatoua villosa EG27
Allaeanthus luzonicus KFC2016
Broussonetia papyrifera SRR1477753
Malaisia scandens EG122
Sloetia elongata Thomas sn
Trilepisium madagascariense Ranktonirina197
Bosqueiopsis gilletii Timberlake5767
Scyphosyce pandurata Thomas6869
Utsetela gabonensis Breteler14096
Dorstenia foetida Burger2844
Treculia africana NZ
Helianthostylis sprucei F16142
T amazonicus GW1522
Trymatococcus oligandrus Hahn3649
Brosimum multinervium Jaramillo574
Brosimum rubescens McPherson20944
Brosimum utile Galdames4403
Brosimum parinarioides Vasquez3817
Brosimum potabile Maciel1552
Brosimum longifolium Vasquez4044 run7
Brosimum lactescens Rivera2204
B lactescens Moza138
Brosimum costaricanum Kernan27
Brosimum acutifolium Lilleen4457
Brosimum alicastrum EG23
Brosimum alicastrum bolivarinese Berg441
Brosimum guianense Morales5258
Brosimum cf guianense Irwin54583
Brosimum gauidichaudii Mendes332
B glaucum Boom13882
91.6
66
56
33.9
23
0
Upper
Cretaceous
Paleocene
Eocene
Oligocene
Paleogene
Miocene
Neogene
Pl
P
Q
(c)
Trema orientale SRR5674478
Maclura pomifera EG139
Artocarpus heterophyllus EG98
Streblus glaber EG78
Parartocarpus venenosus NZ874
Antiaropsis descipiens NZ281
Ficus macrophylla EG30
Fatoua villosa EG27
Allaeanthus luzonicus KFC2016
Broussonetia papyrifera SRR1477753
Malaisia scandens EG122
Sloetia elongata Thomas sn
Trilepisium madagascariense Ranktonirina197
Bosqueiopsis gilletii Timberlake5767
Scyphosyce pandurata Thomas6869
Utsetela gabonensis Breteler14096
Dorstenia foetida Burger2844
Treculia africana NZ
Helianthostylis sprucei F16142
T amazonicus GW1522
Trymatococcus oligandrus Hahn3649
Brosimum multinervium Jaramillo574
Brosimum rubescens McPherson20944
Brosimum utile Galdames4403
Brosimum parinarioides Vasquez3817
Brosimum potabile Maciel1552
Brosimum longifolium Vasquez4044 run7
Brosimum lactescens Rivera2204
B lactescens Moza138
Brosimum costaricanum Kernan27
Brosimum acutifolium Lilleen4457
Brosimum alicastrum EG23
Brosimum alicastrum bolivarinese Berg441
Brosimum guianense Morales5258
Brosimum cf guianense Irwin54583
Brosimum gauidichaudii Mendes332
B glaucum Boom13882
91.6
66
56
33.9
23
0
Upper
Cretaceous
Paleocene
Eocene
Oligocene
Paleogene
Miocene
Neogene
Pl
P
Q

Figure S2. Ancestral state reconstructions of ten characters using maximum likelihood under the "ER" model. (A) stipules fully amplexicaulous (0, 1); (B) longer stipules $\leq$ 15 mm (0), > 15 mm (1); (C) lateral veins loop-connected close (0) or far (1) from margin; (D) breeding system monoecious (0), dioecious (1), androdioecious (2); (E) interfloral bracts peltate (0/1); (F) inflorescences unisexual (0), bisexual (1); (G) Staminate perianth well developed (1) or vestigial / lacking (0); (H) pistillode absent (0), present (1); (I) pistillate inflorescence shape globose to ellipsoid (0) or turbinate, cylindrical, or hemispherical (1); (J) pistillate inflorescence surface with notable protuberences (1) or smooth (0); (K) Bracts greater than 1.5 mm (0, 1); (L) pistillate flowers solitary (0) or multiple (1); (M) stigma equal or shorter than style (0), longer than  style (1); (N) stigma disposition angle from vertical: under 49 (0), 45-90 (1), over 90 (2); (O) stigma weakly curved (0), sigmoid; (P) cotyledons unequal (1) or not (0).

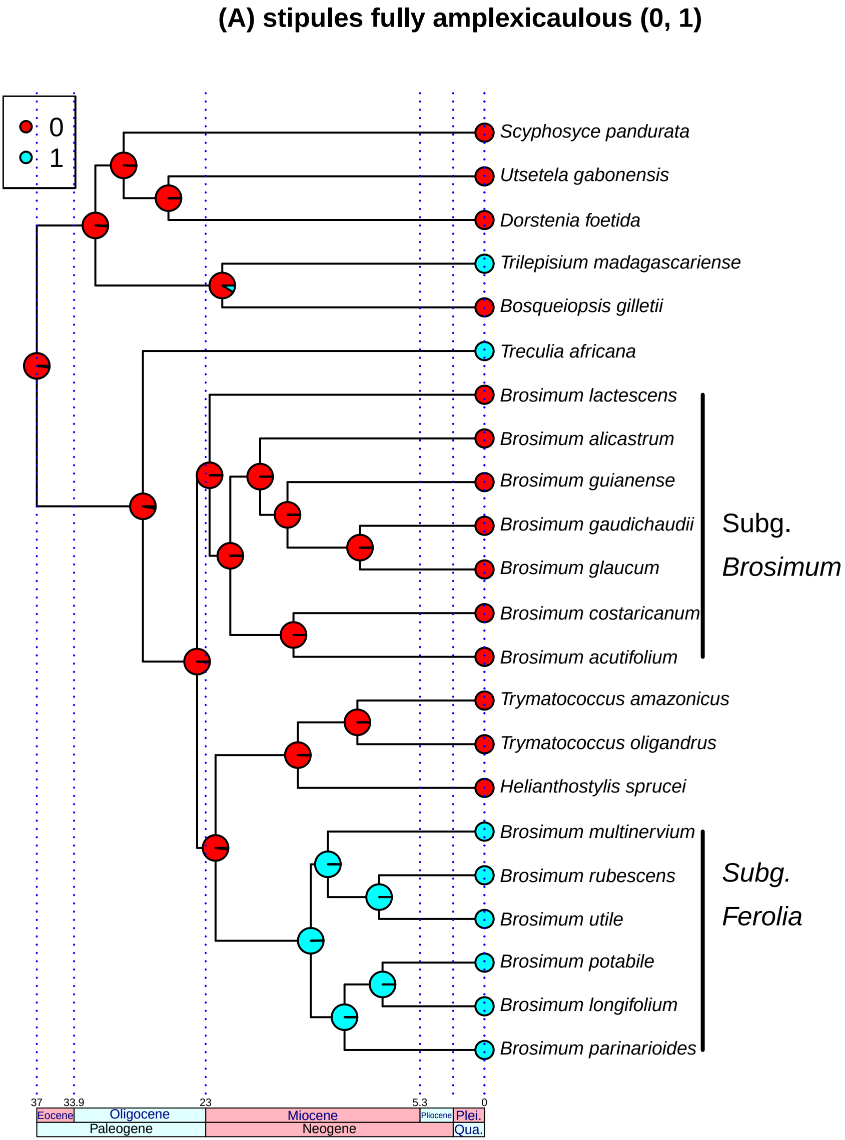

(A) stipules fully amplexicaulous (0, 1)
0
1
Scyphosyce pandurata
Utsetela gabonensis
Dorstenia foetida
Trilepisium madagascariense
Bosqueiopsis gilletii
Treculia africana
Brosimum lactescens
Brosimum alicastrum
Brosimum guianense
Brosimum gaudichaudii
Brosimum glaucum
Brosimum costaricanum
Brosimum acutifolium
Trymatococcus amazonicus
Trymatococcus oligandrus
Helianthostylis sprucei
Brosimum multinervium
Brosimum rubescens
Brosimum utile
Brosimum potabile
Brosimum longifolium
Brosimum parinarioides
Subg. Brosimum
Subg. Ferolia
37
33.9
23
5.3
0
Eocene
Oligocene
Miocene
Pliocene
Plei.
Paleogene
Neogene
Qua.

**(B) longer stipules ... 15 mm (0), > 15 mm (1)**

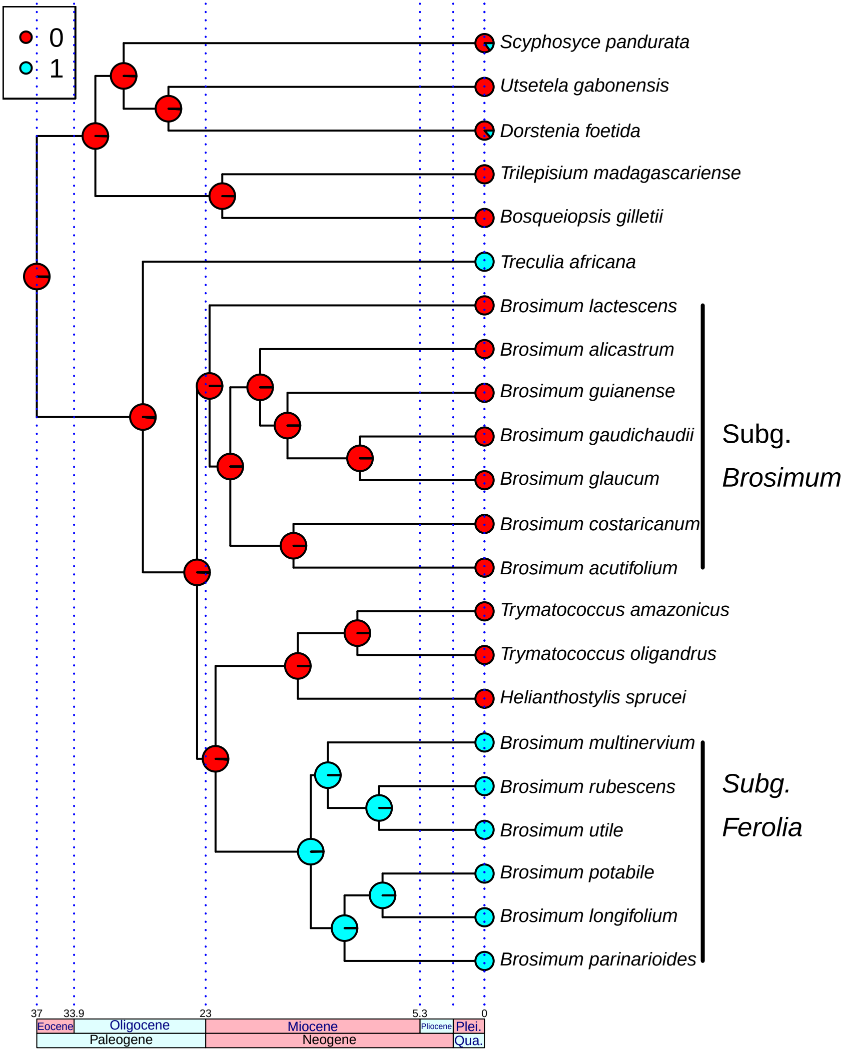

#### (C) lateral veins loop−connected close (0) or far (1) from margin

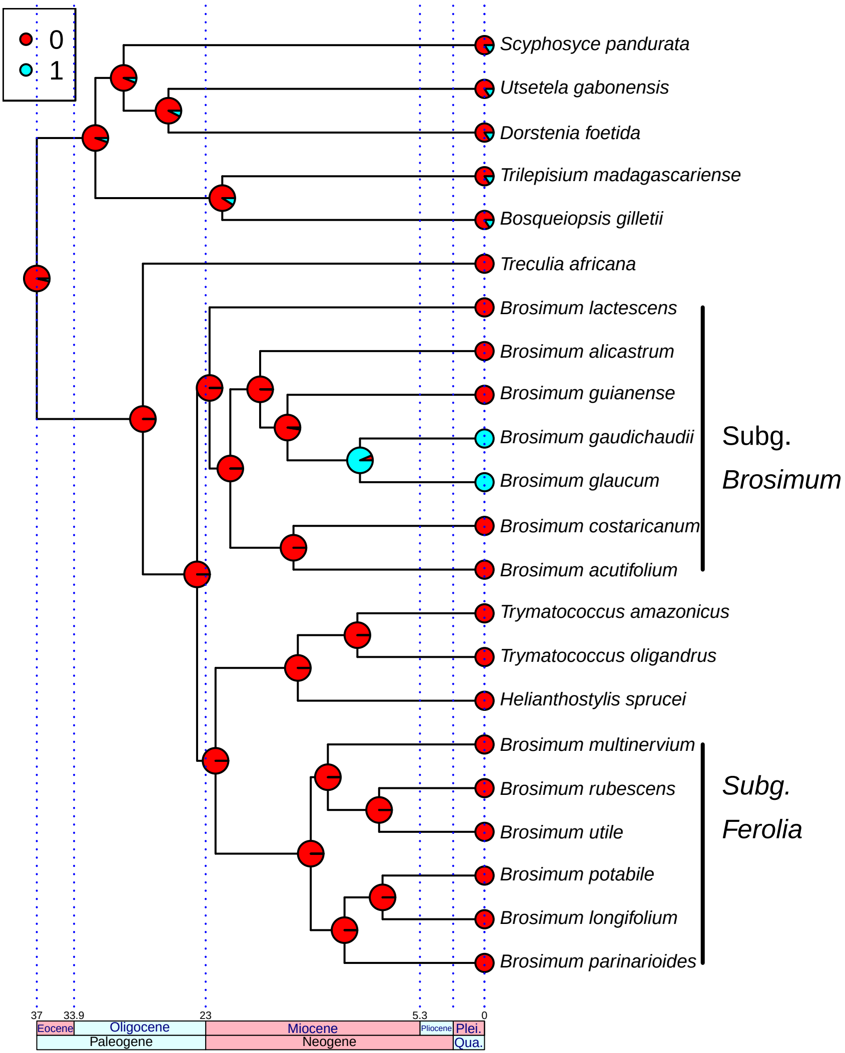

**(D) breeding system monoecious (0), dioecious (1), androdioecious (2)**

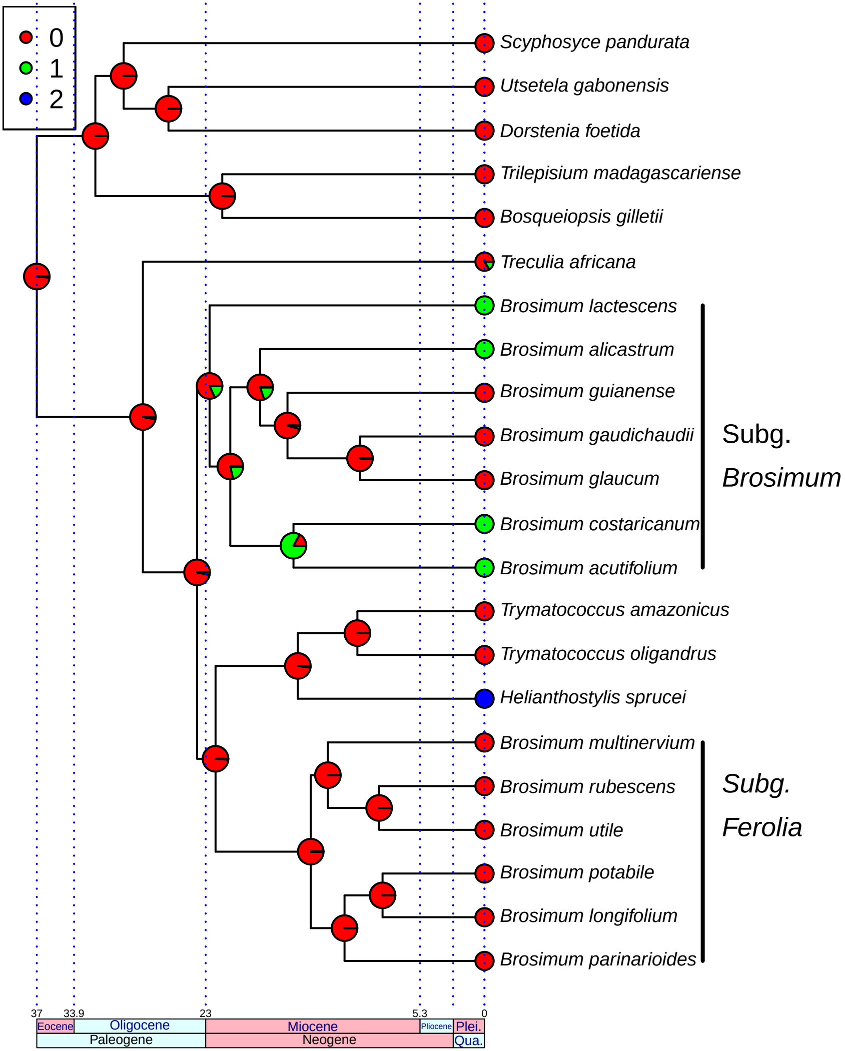

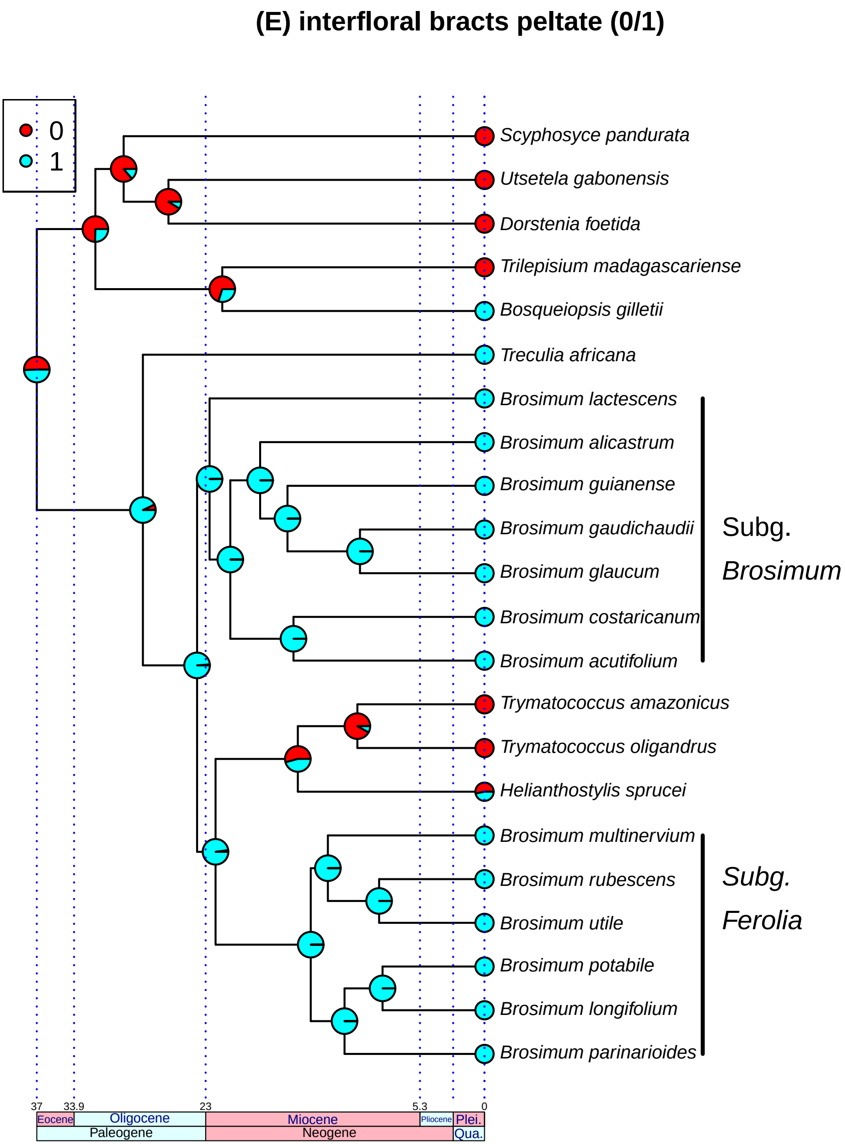

(E) interfloral bracts peltate (0/1)
0
1
Scyphosyce pandurata
Utsetela gabonensis
Dorstenia foetida
Trilepisium madagascariense
Bosqueiopsis gilletii
Treculia africana
Brosimum lactescens
Brosimum alicastrum
Brosimum guianense
Brosimum gaudichaudii
Brosimum glaucum
Brosimum costaricanum
Brosimum acutifolium
Subg. Brosimum
Trymatococcus amazonicus
Trymatococcus oligandrus
Helianthostylis sprucei
Brosimum multinervium
Brosimum rubescens
Brosimum utile
Brosimum potabile
Brosimum longifolium
Brosimum parinarioides
Subg. Ferolia
37
33.9
23
5.3
0
Eocene
Oligocene
Miocene
Pliocene
Plei.
Paleogene
Neogene
Qua.

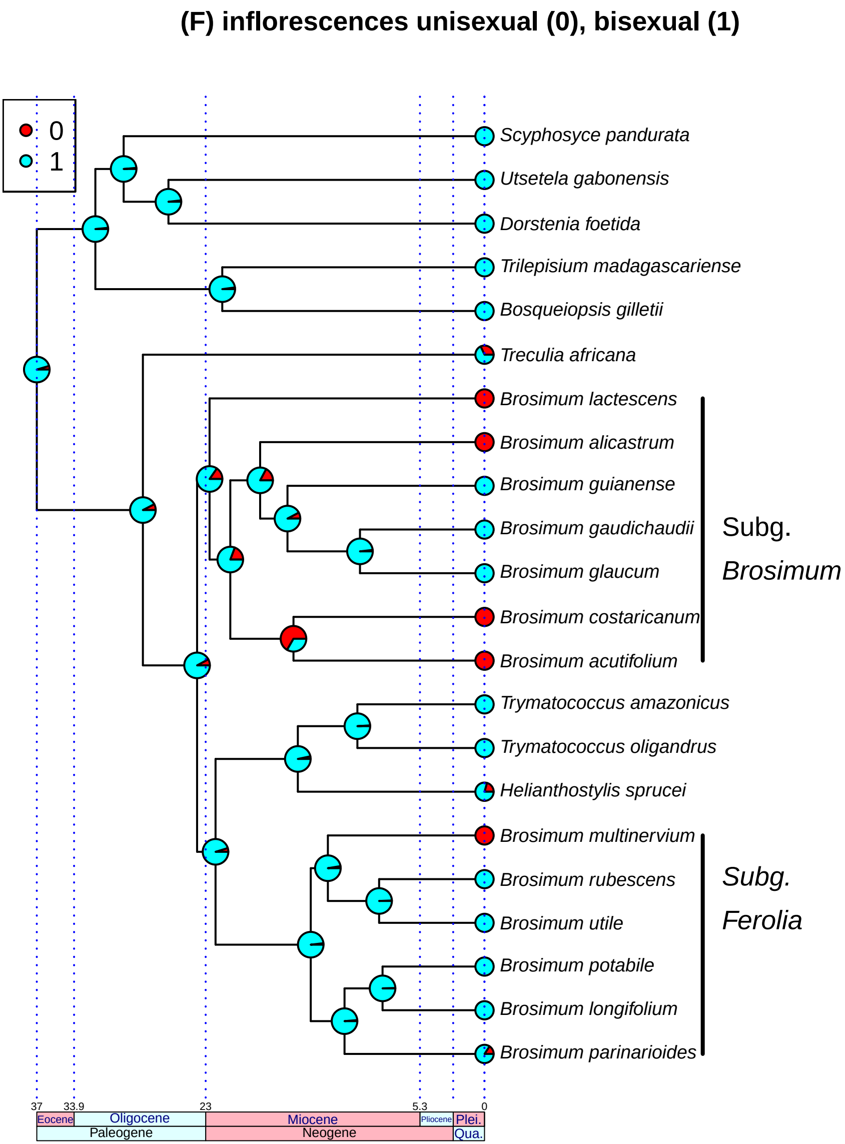
(F) inflorescences unisexual (0), bisexual (1)
0
1
Scyphosyce pandurata
Utsetela gabonensis
Dorstenia foetida
Trilepisium madagascariense
Bosqueiopsis gilletii
Treculia africana
Brosimum lactescens
Brosimum alicastrum
Brosimum guianense
Brosimum gaudichaudii
Brosimum glaucum
Brosimum costaricanum
Brosimum acutifolium
Trymatococcus amazonicus
Trymatococcus oligandrus
Helianthostylis sprucei
Brosimum multinervium
Brosimum rubescens
Brosimum utile
Brosimum potabile
Brosimum longifolium
Brosimum parinarioides
Subg. Brosimum
Subg. Ferolia
37
33.9
23
5.3
0
Eocene
Oligocene
Miocene
Pliocene
Plei.
Paleogene
Neogene
Qua.

#### (G) Staminate perianth well developed (1) or vestigial / lacking (0)

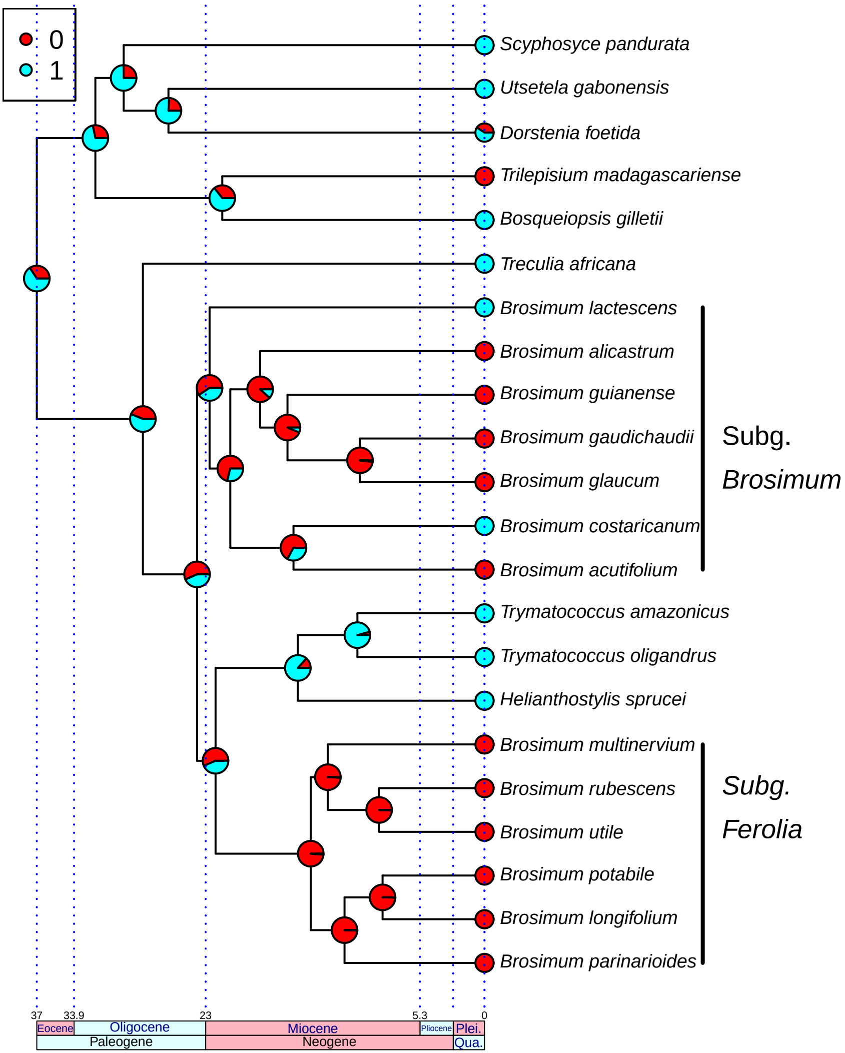

#### (H) pistillode absent (0), present (1)

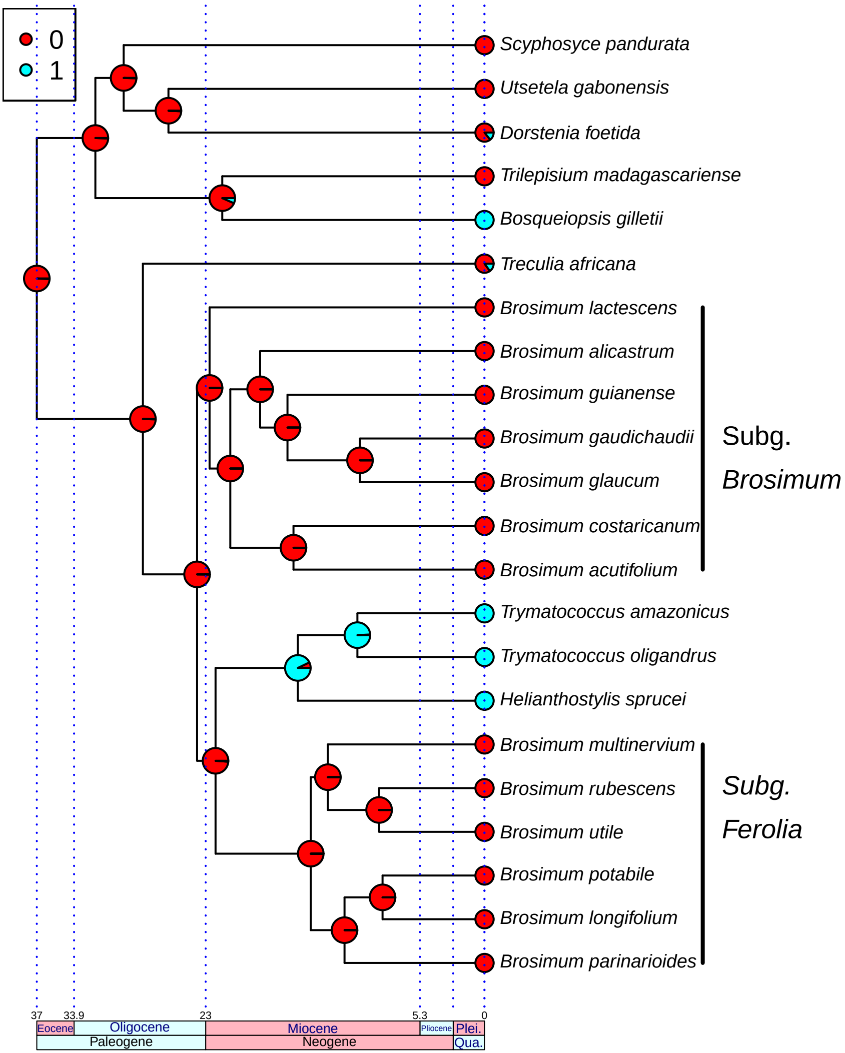

### (I) pistillate inflorescence shape globose to ellipsoid (0) or turbinate, cylindrical, or hemispherical (1)

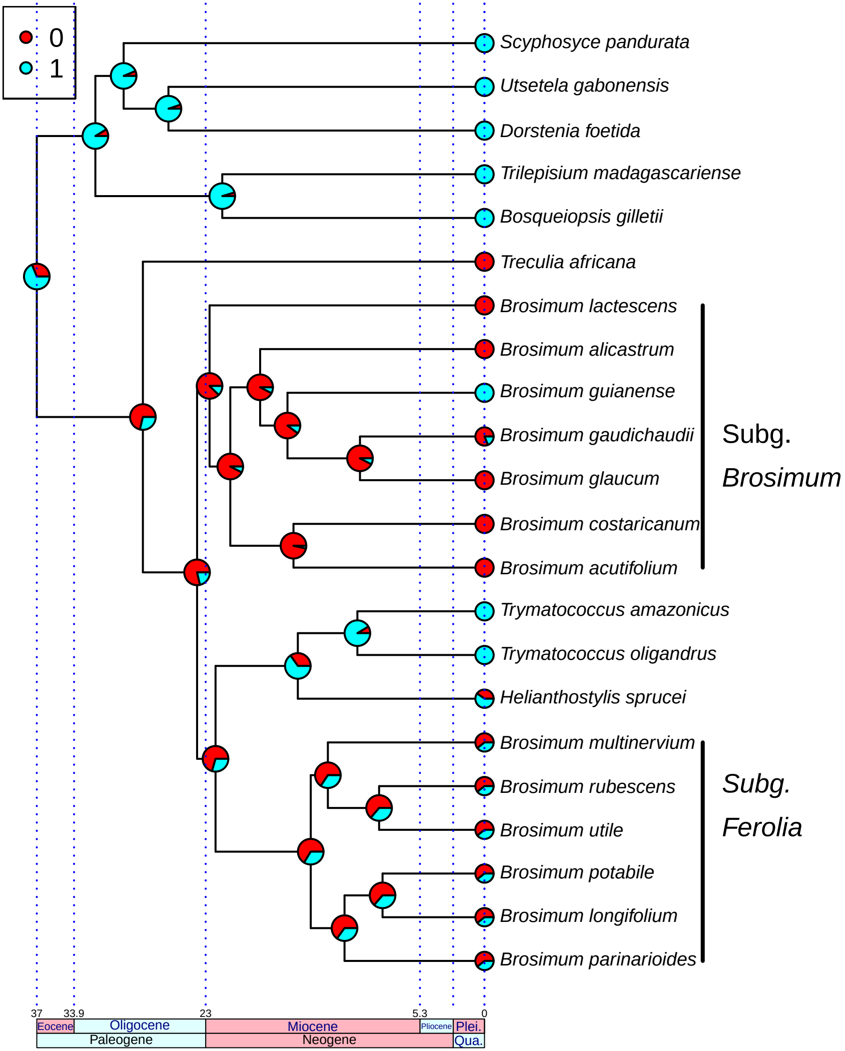

#### (J) pistillate inflorescence surface with notable protuberences (1) or smooth (0)

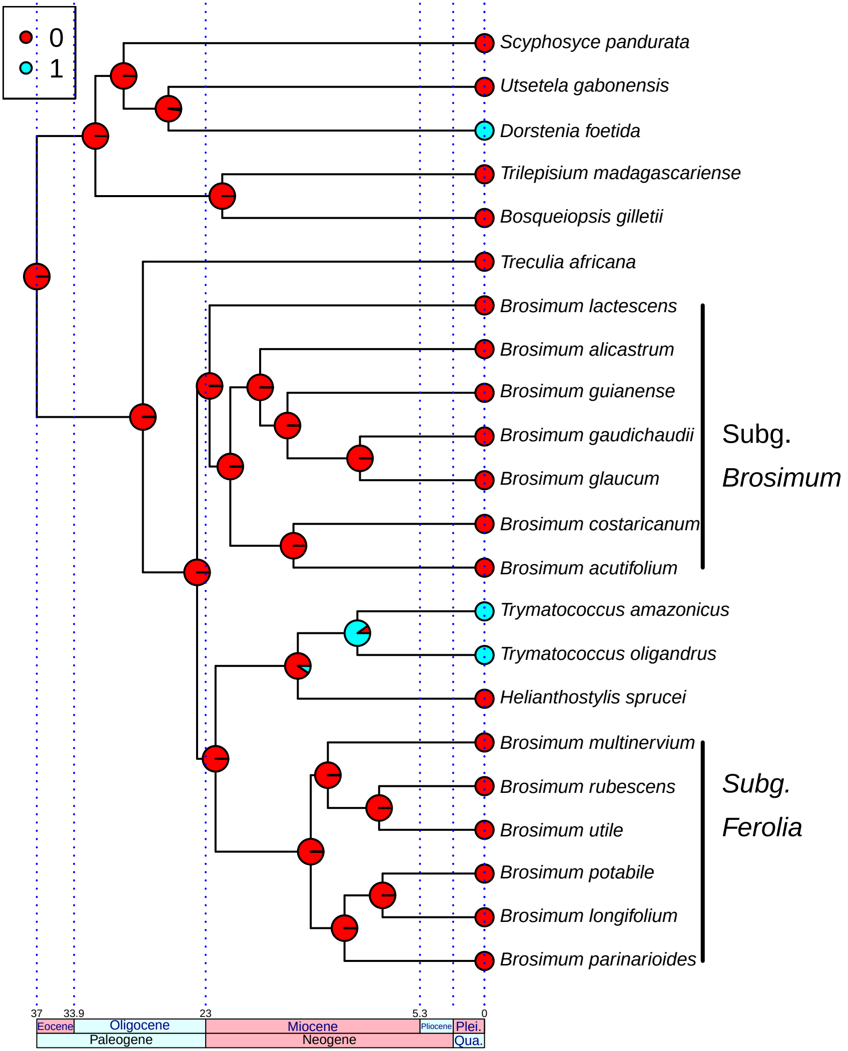

### (K) Bracts greater than 1.5 mm (0, 1)

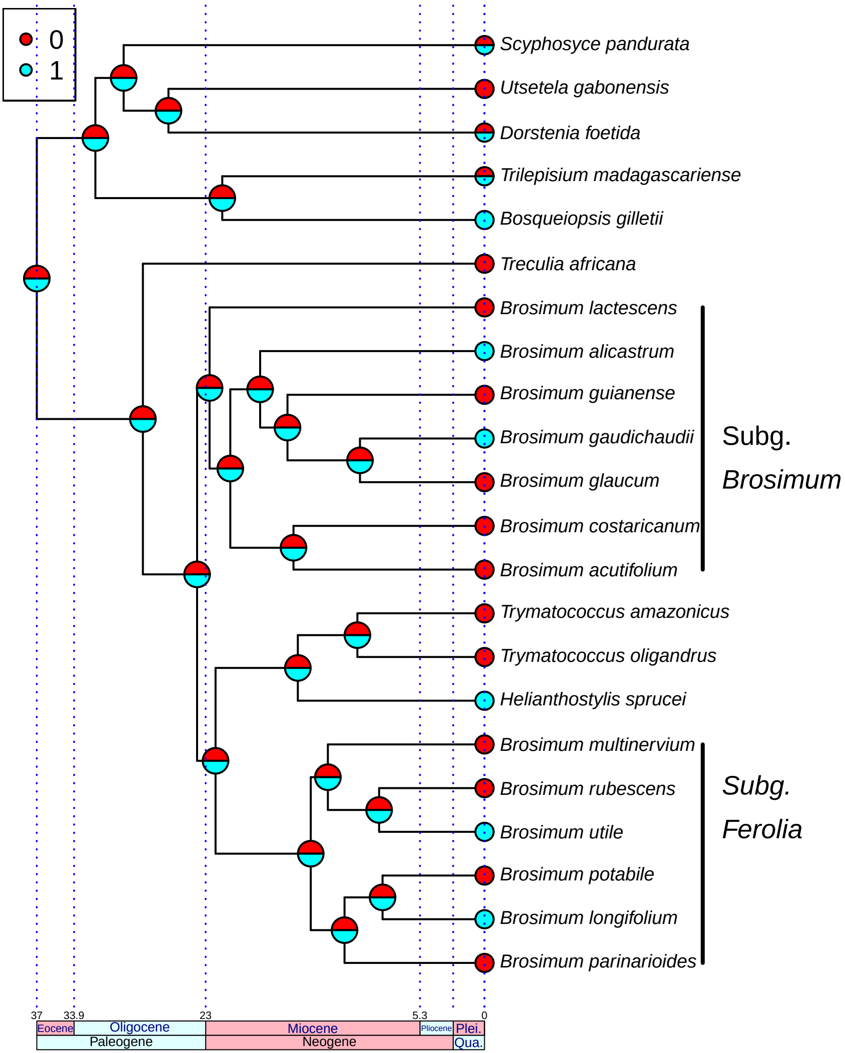

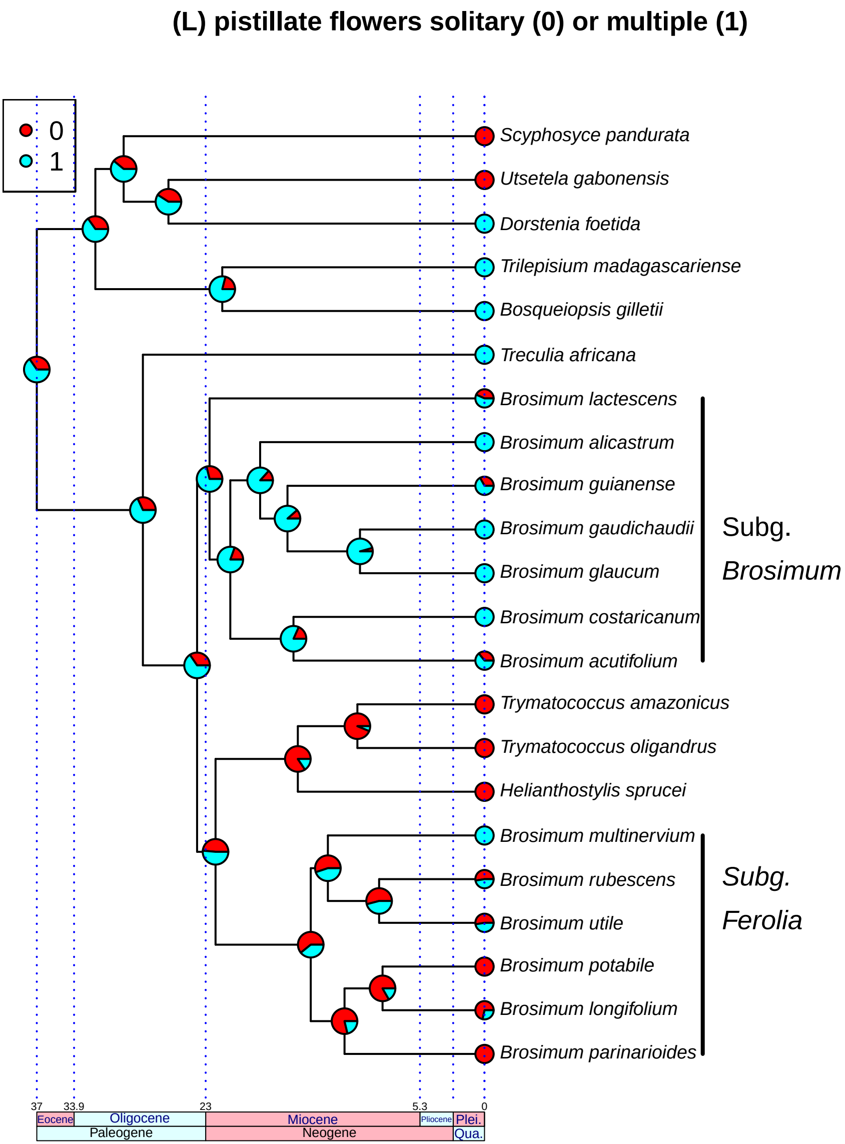
(L) pistillate flowers solitary (0) or multiple (1)
0
1
Scyphosyce pandurata
Utsetela gabonensis
Dorstenia foetida
Trilepisium madagascariense
Bosqueiopsis gilletii
Treculia africana
Brosimum lactescens
Brosimum alicastrum
Brosimum guianense
Brosimum gaudichaudii
Brosimum glaucum
Brosimum costaricanum
Brosimum acutifolium
Trymatococcus amazonicus
Trymatococcus oligandrus
Helianthostylis sprucei
Brosimum multinervium
Brosimum rubescens
Brosimum utile
Brosimum potabile
Brosimum longifolium
Brosimum parinarioides
Subg. Brosimum
Subg. Ferolia
37
33.9
23
5.3
0
Eocene
Oligocene
Miocene
Pliocene
Plei.
Paleogene
Neogene
Qua.

### (M) stigma equal or shorter than style (0), longer than style (1)

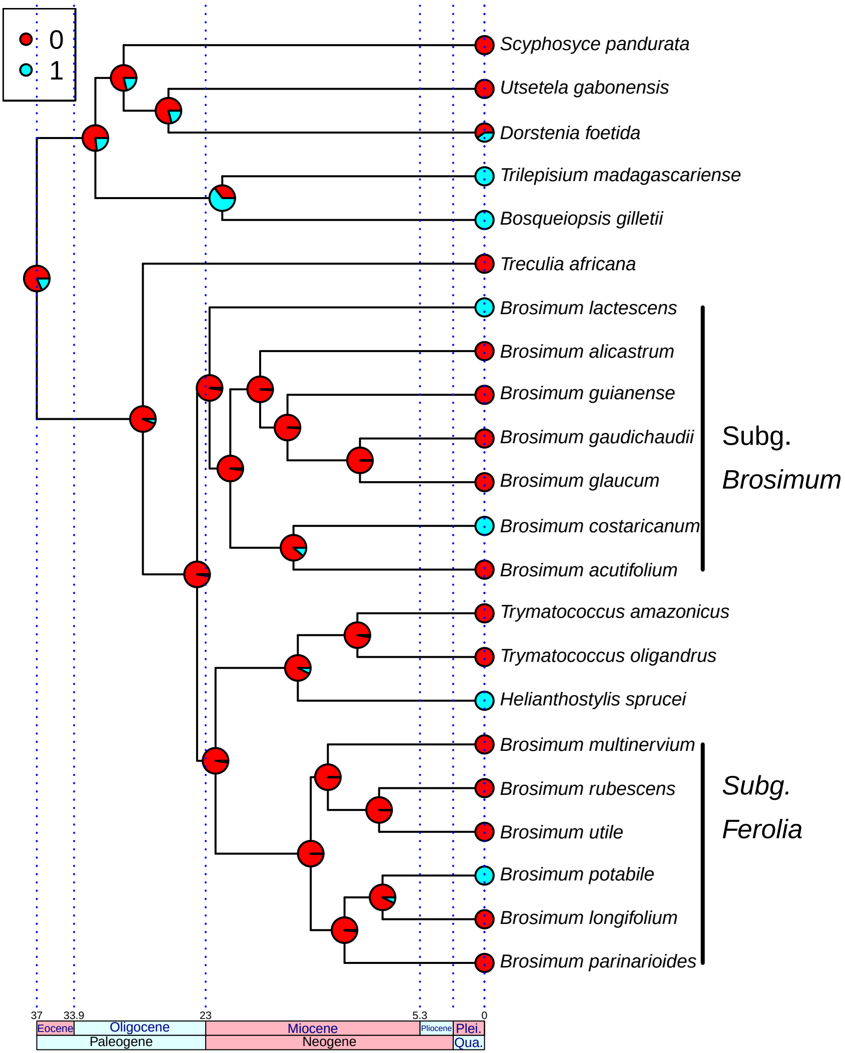

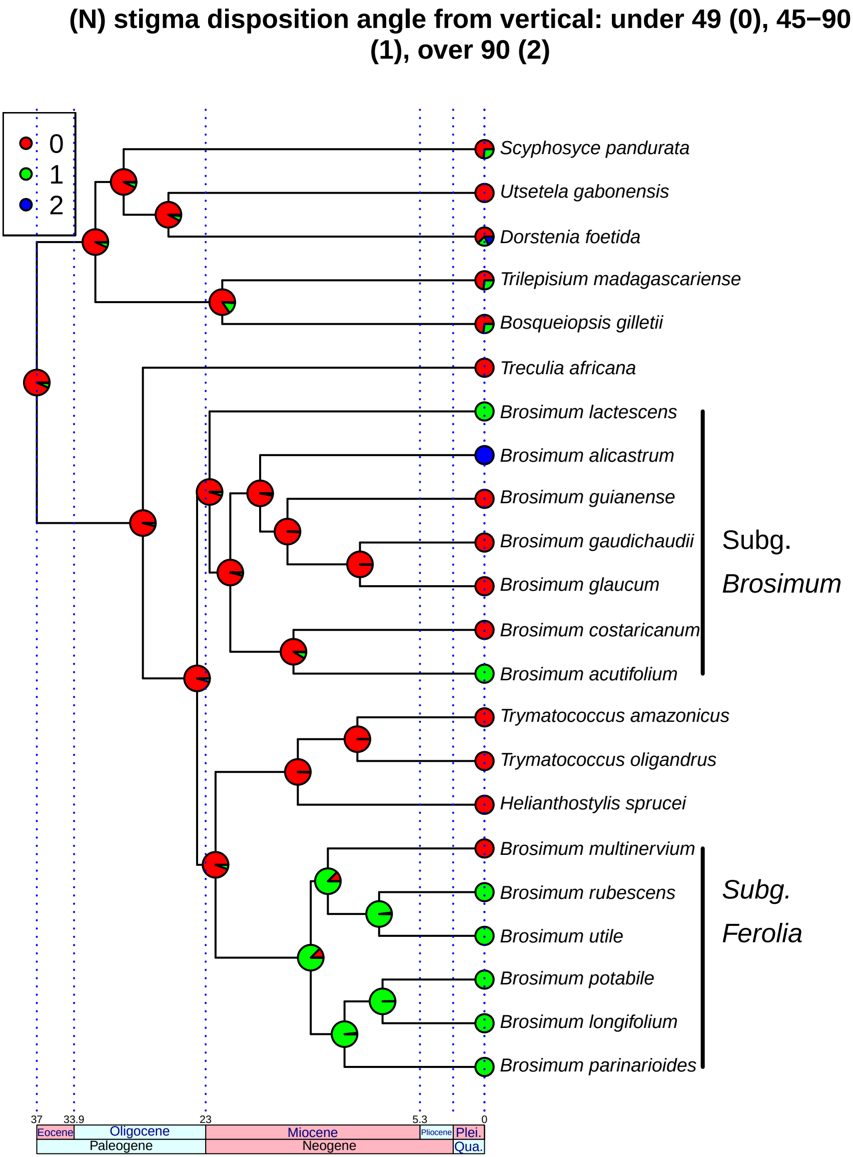
(N) stigma disposition angle from vertical: under 49 (0), 45−90 (1), over 90 (2)
0
1
2
Scyphosyce pandurata
Utsetela gabonensis
Dorstenia foetida
Trilepisium madagascariense
Bosqueiopsis gilletii
Treculia africana
Brosimum lactescens
Brosimum alicastrum
Brosimum guianense
Brosimum gaudichaudii
Brosimum glaucum
Brosimum costaricanum
Brosimum acutifolium
Trymatococcus amazonicus
Trymatococcus oligandrus
Helianthostylis sprucei
Brosimum multinervium
Brosimum rubescens
Brosimum utile
Brosimum potabile
Brosimum longifolium
Brosimum parinarioides
Subg. Brosimum
Subg. Ferolia
37
33.9
23
5.3
0
Eocene
Oligocene
Miocene
Pliocene
Plei.
Paleogene
Neogene
Qua.

### (O) stigma weakly curved (0), sigmoid

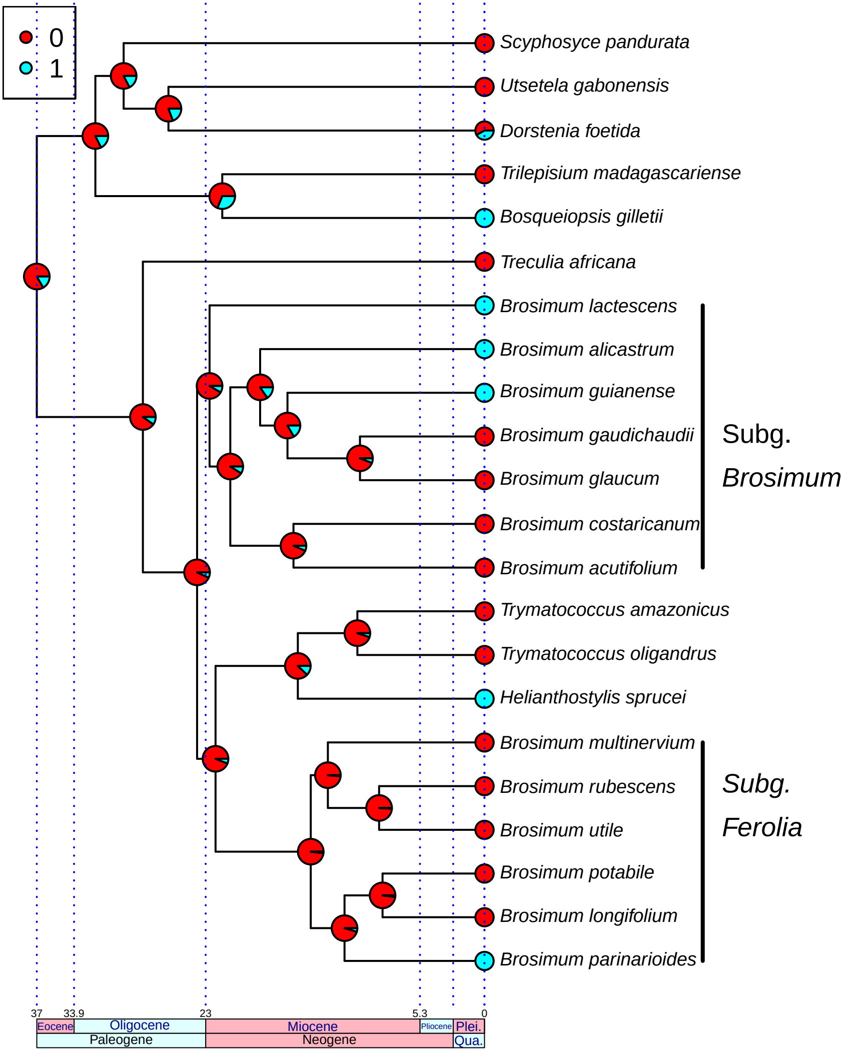

### (P) cotyledons unequal (1) or not (0)

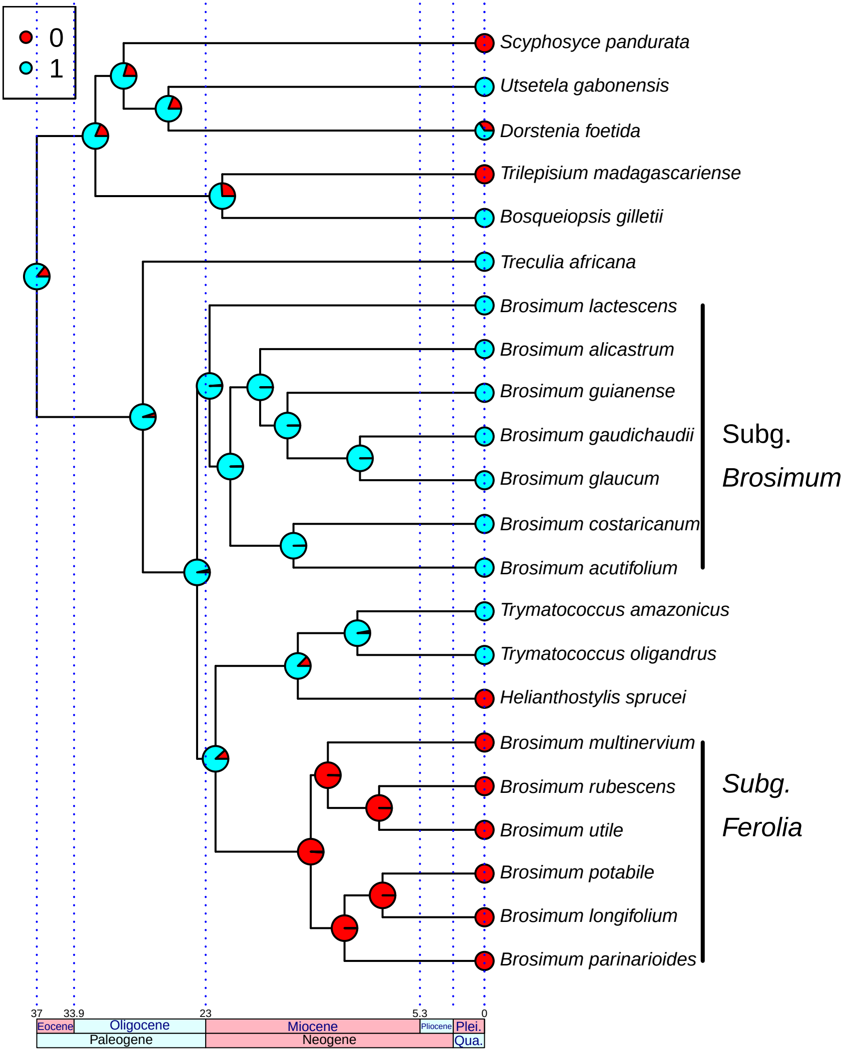

Figure S3. Results of DEC analysis using the biogeographic areas proposed by Antonelli et al. (2018). Left: nodes labeled with the most likely area(s). Right: nodes labels with pie charts indicating relative likelihoods of different scenarios. Areas are AMA (Amazonia), ATF (Atlantic Forests), AGL (Andean Grasslands), CAA (Caatinga), CEC (Cerrado and Chaco), DNO (Dry Northern South America), MES (Mesoamerica), and WIN (West Indies).

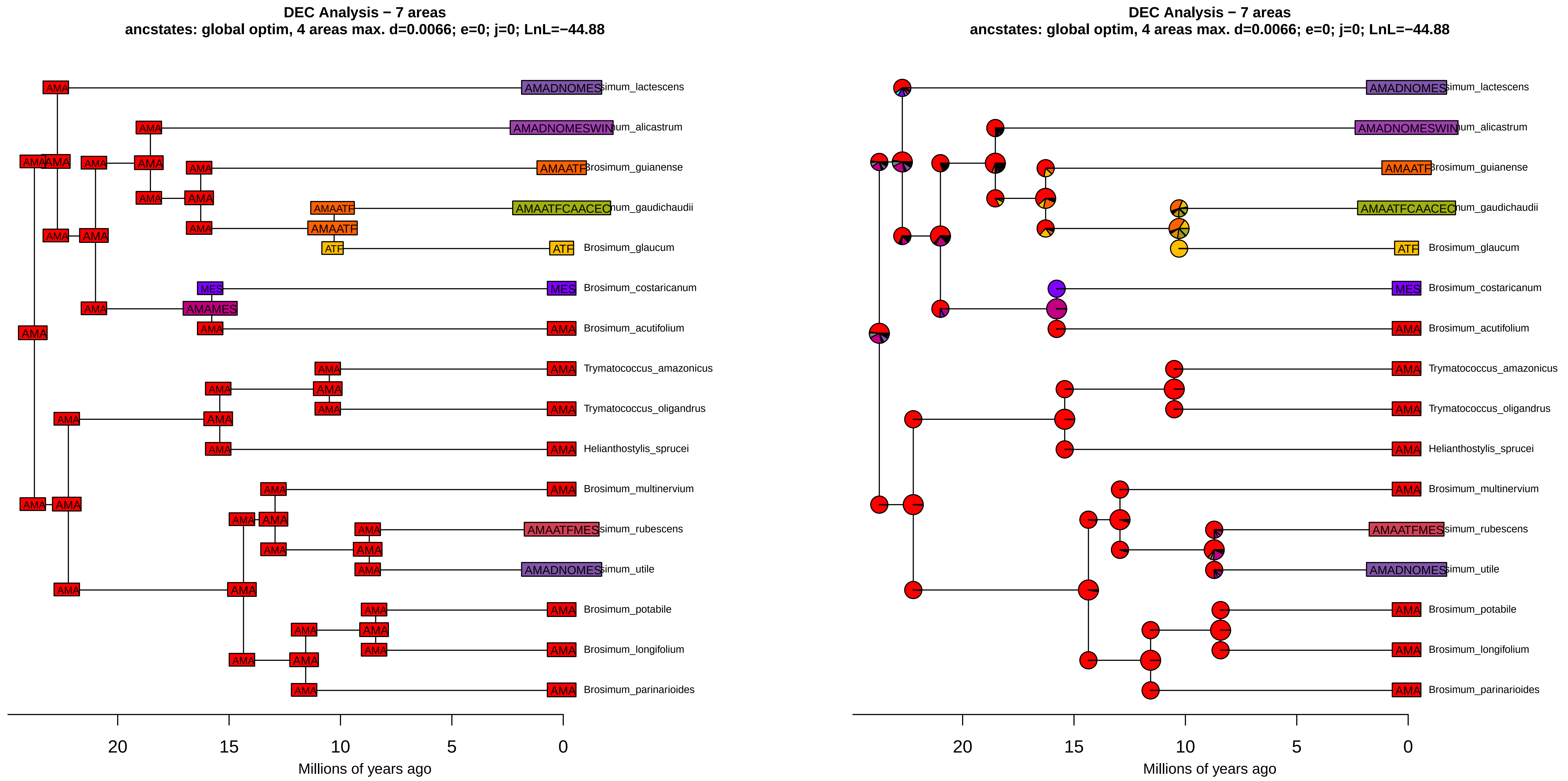
DEC Analysis − 7 areas
ancstates: global optim, 4 areas max. d=0.0066; e=0; j=0; LnL=−44.88
AMADNOMES simum_lactescens
AMADNOMESWIN um_alicastrum
AMAATF Brosimum_guianense
AMAATFCAACEC hum_gaudichaudii
ATF Brosimum_glaucum
MES Brosimum_costaricanum
AMA Brosimum_acutifolium
AMA Trymatococcus_amazonicus
AMA Trymatococcus_oligandrus
AMA Helianthostylis_sprucei
AMA Brosimum_multinervium
AMAATFMES simum_rubescens
AMADNOMES simum_utile
AMA Brosimum_potabile
AMA Brosimum_longifolium
AMA Brosimum_parinarioides
AMAMES
20
15
10
5
0
Millions of years ago
DEC Analysis − 7 areas
ancstates: global optim, 4 areas max. d=0.0066; e=0; j=0; LnL=−44.88
Millions of years ago
