## Appendices: 1A. Character matrix; 1B Area matrix; 1C Anc. recons. model testing. Appendix 2: Analysis workflow scripts for "Phylogenomics of *Brosimum* Sw. (Moraceae) and allied genera, including a revised subgeneric system": Brosimum_Appendix2A.txt

# Appendix 2 to Gardner et al., Phylogenomics of Brosimum and allied genera, including a revised subgeneric system
# Annotated workflow for analyses
# If you find this workflow helpful, please consider citing our paper.

############################################
######### A. Phylogenetic analyses #########
######### Elliot Gardner           #########
############################################

### Required programs:
# GNU Parellel --- https://www.gnu.org/software/parallel/
# Trimmomatic --- http://www.usadellab.org/cms/?page=trimmomatic
# HybPiper (and dependencies) --- https://github.com/mossmatters/HybPiper
# MAFFT --- https://mafft.cbrc.jp/alignment/software/
# TrimAl --- https://github.com/scapella/trimal
# RAxML --- https://github.com/stamatak/standard-RAxML
# ASTRAL --- https://github.com/smirarab/ASTRAL
# fastaselect.tcl --- https://github.com/bioh4x/NeatFreq/tree/master/lib/mira_3.1.15_dev_linux-gnu_x86_64_static/scripts


### Required files:
# Reads in .fastq format (no need to decompress if gzipped)
# "nametable.txt" - a table with two columns; the first containing the read file prefixes and the second contains the desired sample names
# "targets.fna" - HybPiper target file (see HybPiper documentation for formatting instructions)
# "genelist.txt" - a list of gene names that matches the HybPiper target file (de-duplicated, without taxon names)
# "trimbylength.sh" - the following script for filtering sequences base on length (remove the first "#" on each line and paste into a file of that name):
#	### trimbylength.sh
#	### remove sequences from multi-FASTA with two cutoffs: proportion of average length (subtracting Ns), and an absolute minimum in bp.
#	### Elliot Gardner - October 2, 2017
#	### usage: bash trimbylength.sh -i [infile.fasta] -o [outfile.fasta] -p [min fraction of avg length, 0-1] -m [min length in bp]
#	### sequence files must be in current working directory
#	
#	while getopts ":i:o:p:m:" opt; do
#	  case $opt in
#	    i) in="$OPTARG"
#	    ;;
#	    o) out="$OPTARG"
#	    ;;
#	    p) per="$OPTARG"
#	    ;;
#	    m) min="$OPTARG"
#	    ;;
#	  esac
#	done
#	
#	#calculate avg length
#	avg=$(fastalength $in | awk '{ total += $1 } END { print total/NR }')
#	
#	sed -e '/^[^>]/s/N//g' $in > $in.noN
#	
#	fastalength $in.noN | awk '$1 > '"$min"' {print ;}' |awk '{print $1/'"$avg"',$2}' | awk '$1 >= '"$per"' {print $2}' > $in.namestokeep
#	fastaselect.tcl -infile $in -name $in.namestokeep -outfile $out
#	rm $in.namestokeep
#	rm $in.noN
#	printf "input file: "$in"\n"
#	printf "average length: "$avg"\n"
#	printf "input sequences: "$(awk '/>/' $in | wc -l)"\n"
#	printf "output sequences: "$(awk '/>/' $out | wc -l)"\n\n"

### 1. TRIM READS ###
while read a b; do java -jar /home/egardner/bin/Trimmomatic-0.36/trimmomatic-0.36.jar PE -phred33 -threads 16 \
	../reads/"$b"_1.fq.gz ../reads/"$b"_2.fq.gz  \
	"$a"_R1_paired.fastq "$a"_R1_unpaired.fastq "$a"_R2_paired.fastq "$a"_R2_unpaired.fastq \
	ILLUMINACLIP:~/bin/Trimmomatic-0.36/adapters/TruSeq3-PE.fa:2:30:10 \
	HEADCROP:3 LEADING:30 TRAILING:25 SLIDINGWINDOW:4:25 MINLEN:20 ; done < ../nametable.txt

### 2. ASSEMBLE READS ###
while read a b; do ~/bin/HybPiper/reads_first.py --bwa -b ../targets.fna --cpu 16 \
	--prefix $a \
	-r ../trimmed/"$a"_R1_paired.fastq ../trimmed/"$a"_R2_paired.fastq \
	--unpaired ../trimmed/"$a"_R1_unpaired.fastq ; \
	python ~/bin/HybPiper/cleanup.py $a ; \
	python ~/bin/HybPiper/intronerate.py --prefix $a ; \
	done < ../nametable.txt

# check number of genes recovered per sample
wc -l ./**/genes_with_seqs.txt
# if desired, re-run reads_first.py with "--cov_cutoff 4" to increase gene recovery for low-read samples

### 3. FILTER, ALIGN, AND TRIM ("exon" sequences) ###

#Gather and filter
mkdir filter; cd filter
parallel -j 16 cat ../assemblies/**/{}/**/sequences/FNA/{}.FNA '>' {}.raw.fna :::: ../genelist.txt
#Replace the above with the following to retrieve "supercontig" sequences instead
#parallel -j 16 cat ../assemblies/**/{}/**/sequences/intron/{}_supercontig.fasta '>' {}.raw.fna :::: ../genelist.txt

#for each gene, remove sequences less than 50bp or less than 25% of the average length 
parallel -j 16 bash ~/bin/scripts/trimbylength.sh -i {}.raw.fna -o {}.filtered.fna -p 0.25 -m 50 :::: ../genelist.txt

#write by-gene and by-sample counts to files
awk '/>/' *.raw.fna | sort | uniq | sed 's/>//g' > ../samplelist.txt
parallel ' printf {}"\t";awk "/{}/" *.raw.fna | wc -l  ' :::: ../samplelist.txt | sort -k2 -nr > raw_counts.txt
parallel ' printf {}"\t";awk "/{}/" *.filtered.fna | wc -l '  :::: ../samplelist.txt | sort -k2 -nr > filtered_counts.txt
parallel ' printf {}"\t";awk "/>/" {}.filtered.fna | wc -l ' :::: ../genelist.txt | sort -k2 -nr > filtered_counts_by_gene.txt
parallel ' printf {}"\t";awk "/>/" {}.raw.fna | wc -l ' :::: ../genelist.txt | sort -k2 -nr > raw_counts_by_gene.txt

#align and trim
mkdir ../align
cd ../align
parallel mafft ../filter/{}.filtered.fna '>' {}.aln :::: ../genelist.txt

#trim columns with more than 75% gaps.
parallel trimal -gt 0.25 -in {}.aln -out {}.trimmed :::: ../genelist.txt

#optional - discard sequences more than 50% divergent from all other sequences.
#while read a ; do trimal -sident -in $a.aln | awk '/Identity for most similar pair-wise sequences matrix/,EOF' | awk '!/#/' | awk '$2>0.5 {print $1}' > $a.names ; fastaselect.tcl -infile $a.aln -outfile $a.aln.cut -name $a.names ; rm $a.names ; done < ../genelist_cut.txt
#parallel trimal -gt 0.25 -in {}.aln.cut -out {}.trimmed :::: ../genelist.txt

#build supermatrix and partition file
python ~/bin/HybPiper/fasta_merge.py --fastafiles *.trimmed --raxml DNA > supermatrix.exons.fasta
mv partition.raxml supermatrix.exons.part
sed -i 's/ <unknown description>//g' supermatrix.exons.fasta
cd ..

### 4. PHYLOGENETIC ANALYSES ###

# Estimate ML tree from supermatrix
mkdir supermatrix; cd supermatrix
mv ../align/supermatrix* .
raxmlHPC-PTHREADS-AVX -T 16 -n supermatrix.exon.out -s supermatrix.exons.fasta -q supermatrix.exons.part -p 12345 -x 12345 -N 500 -c 25 -f a -m GTRGAMMA -o Trema_orientale_SRR5674478

# Gene trees
#generate gene trees with RAxML
mkdir ../genetrees; cd ../genetrees
parallel -j 6 raxmlHPC-PTHREADS-AVX -T 2 -n {}.out -s ../align/{}.trimmed -p 12345 -x 12345 -N 200 -c 25 -f a -m GTRGAMMA :::: ../genelist_cut.txt
#sort files
mkdir info; mv RAxML_info.* ./info/
mkdir bipartitionsBranchLabels; mv RAxML_bipartitionsBranchLabels.* ./bipartitionsBranchLabels/
mkdir bipartitions; mv RAxML_bipartitions.* ./bipartitions/
mkdir bestTree; mv RAxML_bestTree.* ./bestTree/
mkdir bootstrap; mv RAxML_bootstrap.* ./bootstrap/
mkdir collapsed
#collapse nodes under 30%
parallel sumtrees.py --replace bootstrap/RAxML_bootstrap.{}.out -f 0.33 -F newick --suppress-annotations -o collapsed/RAxML_collapsed.{}.tre -i newick -t bestTree/RAxML_bestTree.{}.out :::: ../genelist.txt
parallel "sed -i 's/\[&U\] //g' collapsed/RAxML_collapsed.{}.tre" :::: ../genenlist.txt
parallel "sed -i \"s/'//g\" collapsed/RAxML_collapsed.{}.tre" :::: ../genelist.txt
cat ./collapsed/RAxML_collapsed.* > bestTrees_collapsed.tre
ls ./bootstrap/* > bootstraplist.txt

# Estimate species tree with ASTRAL
# with LPP (quartet scores)
java -jar ~/bin/ASTRAL/astral.5.6.1.jar -i bestTrees_collapsed.tre -t 1 -o astral.collapsed30.lpp.tre
# with bootstrap
java -jar ~/bin/ASTRAL/astral.5.6.1.jar -i bestTrees_collapsed.tre -b bootstraplist.txt -g -r 150 -o astral.collapsed30.100bs.tre
