## Appendices: 1A. Character matrix; 1B Area matrix; 1C Anc. recons. model testing. Appendix 2: Analysis workflow scripts for "Phylogenomics of *Brosimum* Sw. (Moraceae) and allied genera, including a revised subgeneric system": Brosimum_Appendix2D.txt

# Appendix 1 to Gardner et al., Phylogenomics of Brosimum and allied genera, including a revised subgeneric system
# Annotated workflow for analyses
# If you find this workflow helpful, please consider citing our paper.

################################################################
######### D. Dispersal-Extinction-Cladogensis Analyses #########
######### Elliot Gardner                               #########
################################################################

### This is an R script.
# Required files:
# baseml_noOG.tre - ultrametric tree with no outgroups
# brosimum_biogeo_4.txt - matrix with 4 areas
# brosimum_biogeo_8.txt - matrix with 8 areas

library(openxlsx)
library(GenSA)
library(FD)
library(snow)
library(parallel)
library(rexpokit)
library(cladoRcpp)
library(BioGeoBEARS)

#set working directory
setwd("PATH/TO/WORKING/DIRECTORY/")

extdata_dir = np(system.file("extdata", package="BioGeoBEARS"))

trfn = "baseml_noOG.tre"
tr = read.tree(trfn)
#use one or the other of these:
#geogfn = "brosimum_biogeo_4.txt"
geogfn = "brosimum_biogeo_10.txt"
tipranges = getranges_from_LagrangePHYLIP(lgdata_fn=geogfn)
max(rowSums(dfnums_to_numeric(tipranges@df)))
max_range_size = 4
#check to make sure this isn't crazy:
numstates_from_numareas(numareas=4, maxareas=3, include_null_range=TRUE)

# verbatim from http://phylo.wikidot.com/biogeobears#script
#######################################################
# Run DEC
#######################################################

# Intitialize a default model (DEC model)
BioGeoBEARS_run_object = define_BioGeoBEARS_run()

# Give BioGeoBEARS the location of the phylogeny Newick file
BioGeoBEARS_run_object$trfn = trfn

# Give BioGeoBEARS the location of the geography text file
BioGeoBEARS_run_object$geogfn = geogfn

# Input the maximum range size
BioGeoBEARS_run_object$max_range_size = max_range_size

BioGeoBEARS_run_object$min_branchlength = 0.000001    # Min to treat tip as a direct ancestor (no speciation event)
BioGeoBEARS_run_object$include_null_range = TRUE    # set to FALSE for e.g. DEC* model, DEC*+J, etc.
# (For DEC* and other "*" models, please cite: Massana, Kathryn A.; Beaulieu, 
#  Jeremy M.; Matzke, Nicholas J.; O’Meara, Brian C. (2015). Non-null Effects of 
#  the Null Range in Biogeographic Models: Exploring Parameter Estimation in the 
#  DEC Model. bioRxiv,  http://biorxiv.org/content/early/2015/09/16/026914 )
# Also: search script on "include_null_range" for other places to change

# Set up a time-stratified analysis:
# 1. Here, un-comment ONLY the files you want to use.
# 2. Also un-comment "BioGeoBEARS_run_object = section_the_tree(...", below.
# 3. For example files see (a) extdata_dir, 
#  or (b) http://phylo.wikidot.com/biogeobears#files
#  and BioGeoBEARS Google Group posts for further hints)
#
# Uncomment files you wish to use in time-stratified analyses:
#BioGeoBEARS_run_object$timesfn = "timeperiods.txt"
#BioGeoBEARS_run_object$dispersal_multipliers_fn = "manual_dispersal_multipliers.txt"
#BioGeoBEARS_run_object$areas_allowed_fn = "areas_allowed.txt"
#BioGeoBEARS_run_object$areas_adjacency_fn = "areas_adjacency.txt"
#BioGeoBEARS_run_object$distsfn = "distances_matrix.txt"
# See notes on the distances model on PhyloWiki's BioGeoBEARS updates page.

# Speed options and multicore processing if desired
BioGeoBEARS_run_object$on_NaN_error = -1e50    # returns very low lnL if parameters produce NaN error (underflow check)
BioGeoBEARS_run_object$speedup = TRUE          # shorcuts to speed ML search; use FALSE if worried (e.g. >3 params)
BioGeoBEARS_run_object$use_optimx = "GenSA"    # if FALSE, use optim() instead of optimx()
BioGeoBEARS_run_object$num_cores_to_use = 1
# (use more cores to speed it up; this requires
# library(parallel) and/or library(snow). The package "parallel" 
# is now default on Macs in R 3.0+, but apparently still 
# has to be typed on some Windows machines. Note: apparently 
# parallel works on Mac command-line R, but not R.app.
# BioGeoBEARS checks for this and resets to 1
# core with R.app)

# Sparse matrix exponentiation is an option for huge numbers of ranges/states (600+)
# I have experimented with sparse matrix exponentiation in EXPOKIT/rexpokit,
# but the results are imprecise and so I haven't explored it further.
# In a Bayesian analysis, it might work OK, but the ML point estimates are
# not identical.
# Also, I have not implemented all functions to work with force_sparse=TRUE.
# Volunteers are welcome to work on it!!
BioGeoBEARS_run_object$force_sparse = FALSE    # force_sparse=TRUE causes pathology & isn't much faster at this scale

# This function loads the dispersal multiplier matrix etc. from the text files into the model object. Required for these to work!
# (It also runs some checks on these inputs for certain errors.)
BioGeoBEARS_run_object = readfiles_BioGeoBEARS_run(BioGeoBEARS_run_object)

# Divide the tree up by timeperiods/strata (uncomment this for stratified analysis)
#BioGeoBEARS_run_object = section_the_tree(inputs=BioGeoBEARS_run_object, make_master_table=TRUE, plot_pieces=FALSE)
# The stratified tree is described in this table:
#BioGeoBEARS_run_object$master_table

# Good default settings to get ancestral states
BioGeoBEARS_run_object$return_condlikes_table = TRUE
BioGeoBEARS_run_object$calc_TTL_loglike_from_condlikes_table = TRUE
BioGeoBEARS_run_object$calc_ancprobs = TRUE    # get ancestral states from optim run

# Set up DEC model
# (nothing to do; defaults)

# Look at the BioGeoBEARS_run_object; it's just a list of settings etc.
BioGeoBEARS_run_object

# This contains the model object
BioGeoBEARS_run_object$BioGeoBEARS_model_object

# This table contains the parameters of the model 
BioGeoBEARS_run_object$BioGeoBEARS_model_object@params_table

# Run this to check inputs. Read the error messages if you get them!
check_BioGeoBEARS_run(BioGeoBEARS_run_object)

# For a slow analysis, run once, then set runslow=FALSE to just 
# load the saved result.
runslow = TRUE
#resfn = "Brosimum_DEC_4areas.Rdata"
resfn = "Brosimum_DEC_8areas.Rdata"
if (runslow)
    {
    res = bears_optim_run(BioGeoBEARS_run_object)
    res    

    save(res, file=resfn)
    resDEC = res
    } else {
    # Loads to "res"
    load(resfn)
    resDEC = res
    }
                

#######################################################
# PDF plots
#######################################################
pdffn = "Brosimum_DEC_unconstrained.pdf"
pdf(pdffn, width=16, height=8)
par(mfcol=c(1,2))

#######################################################
# Plot ancestral states - DEC
#######################################################
analysis_titletxt ="Brosimum DEC Analysis"

# Setup
results_object = resDEC
scriptdir = np(system.file("extdata/a_scripts", package="BioGeoBEARS"))

#plot them on the same page

# States
res2 = plot_BioGeoBEARS_results(results_object, analysis_titletxt, addl_params=list("j"), plotwhat="text", label.offset=0.45, tipcex=0.6, statecex=0.7, splitcex=0.6, titlecex=0.8, plotsplits=TRUE, cornercoords_loc=scriptdir, include_null_range=TRUE, tr=tr, tipranges=tipranges)

# Pie chart
plot_BioGeoBEARS_results(results_object, analysis_titletxt, addl_params=list("j"), plotwhat="pie", label.offset=0.45, tipcex=0.6, statecex=0.7, splitcex=0.6, titlecex=0.8, plotsplits=TRUE, cornercoords_loc=scriptdir, include_null_range=TRUE, tr=tr, tipranges=tipranges)

dev.off()  # Turn off PDF
cmdstr = paste("open ", pdffn, sep="")
system(cmdstr) # Plot it
